# Biochemical characterization of *Bacillus anthracis* sortase B: Use in sortase-mediated ligation and substrate recognition dependent on residues beyond the canonical pentapeptide binding motif for sortase enzymes

**DOI:** 10.1101/2024.05.07.593036

**Authors:** Sophie N. Jackson, Jadon M. Blount, Kayla A. Croney, Darren E. Lee, Justin W. Ibershof, Kyle M. Whitham, James McCarty, John M. Antos, Jeanine F. Amacher

## Abstract

Sortases are cysteine transpeptidases located on the surface of Gram-positive bacteria. These critical enzymes facilitate the attachment of proteins to the cell wall, and are potential targets for novel antibiotic development, as well as versatile tools in protein engineering applications. Although there are six classes of sortases recognized, class A sortases (SrtA) are the most widely studied and utilized. SrtA enzymes recognize the canonical Cell Wall Sorting Signal (CWSS), LPXTG, where X=any amino acid, although work in recent years identified additional promiscuity in multiple positions of this recognition motif. Much less is known about Class B sortases (SrtB), which target a distinct sequence, typically with an N-terminal Asn, e.g., variations of NPXTG or NPQTN. Although understudied overall, two SrtB enzymes have previously been shown to be specific for heme transporter proteins, and *in vitro* experiments with the catalytic domains of these enzymes reveal activities significantly worse than SrtA from the same organisms. Here, we use protein biochemistry, structural analyses, and computational simulations to better understand and characterize these enzymes, specifically investigating *Bacillus anthracis* SrtB (baSrtB) as a model SrtB protein. Structural modeling predicts a plausible enzyme-substrate complex, which is verified by mutagenesis of binding cleft residues at several positions. Furthermore, residues N- and C-terminal to the pentapeptide recognition motif are critical for observed activity. We also use chimeric proteins to identify a single site that improves baSrtB activity by ∼4-fold and use purified protein substrates to validate sortase-mediated ligation of two proteins using SrtB enzymes for the first time. Taken together, these studies provide insight into SrtB-target binding as well as evidence that SrtB enzymes can be modified to be of potential use in protein engineering.

## Introduction

Surface proteins enable bacteria to perform functions that are critical for survival. In pathogens, these proteins are frequently virulence factors, although they also include those for environmental sensing and other cellular functions.^1,2^ Many surface proteins in Gram-positive bacteria are anchored by sortase enzymes.^1–3^ Sortases are membrane-anchored cysteine transpeptidases that attach target proteins to the cell wall via a ping-pong ligation reaction.^1–3^ The first sortase enzyme discovered was the class A sortase from *Staphylococcus aureus* (saSrtA).^3,4^ Due to their critical role in the bacterial life cycle, sortase enzymes have emerged as potential targets for novel antibiotic development.^5–7^ Furthermore, because of their ability to ligate two protein-derived sequences together, sortases are powerful protein engineering tools, via sortase-mediated ligation (SML) or *sortagging* applications.^8–11^ SaSrtA remains the most widely utilized enzyme in SML experiments, as well as the best studied sortase. However, >10,000 sortase sequences have been identified from over 1,000 bacterial species, which are generally grouped into six classes, A-F, based on primary sequence.^12–14^

The sortase catalytic mechanism proceeds in a number of distinct steps via the action of a conserved triad of residues (His-Cys-Arg), with recent work also suggesting a key role for a conserved Thr residue immediately preceding the active site Cys.^1,15–18^ Class A sortases generally recognize and anchor proteins containing a LPXTG motif within their Cell Wall Sorting Signal (CWSS), where X=any amino acid and Leu=P4, Pro=P3, X=P2, T=P1, and G=P1’.^2,19^ Upon substrate binding by the sortase enzyme, the active site Cys nucleophile cleaves between the P1/P1’ positions, resulting in an acyl-enzyme intermediate (LPXT-enzyme). To facilitate formation of this intermediate, it has been proposed that the side chain hydroxyl of the highly conserved Thr residue, as well as the backbone amides of the catalytic Cys and the residue immediately C-terminal to the conserved His, serve as an oxyanion hole to stabilize charged intermediates en route to the acyl-enzyme state.^15,17,20,21^ Next, the acyl-enzyme is resolved by nucleophilic attack by the amine terminus (often Gly or Ala) of a second enzyme substrate, leading to release of the final ligated product, LPXT-G/A.^1^ Throughout this entire process, the conserved Arg likely serves to bind and position the LPXT fragment, primarily through polar contacts with the carbonyls of the P4 Leu and P3 Pro.^15,17,20^ Previous work also suggested a possible role for this Arg in stabilizing oxyanion intermediates.^1^ Sortases from all classes studied to date appear to follow a similar mechanism, with differing classes regulated by distinct structural features of the sortase enzyme and/or variations in the substrate sequence(s) recognized.^2,12^

While class A sortases are often referred to as general housekeeping enzymes, class B sortases (SrtBs) are more specific. Identified SrtB enzymes include those that recognize a specific heme transporter protein, IsdC, from the iron-regulated surface determinant (Isd) family.^22,23^ IsdC is the only known substrate for *S. aureus* and *Bacillus anthracis* SrtB, which recognize NPQTN and NPKTG sequences, respectively.^24-27^ Other studied SrtB enzymes are involved in different processes, e.g., *Streptococcus pyogenes* SrtB polymerizes the pilus and requires a cofactor, SipA, *Clostridioides difficile* SrtB is suspected to recognize several proteins involved in adhesion, including CD0386, and cell wall hydrolysis, and *Listeria monocytogenes* SrtB targets proteins likely involved in iron acquisition, Lmo2185 and Lmo2186, but which utilize a different pathway than *S. aureus* or *B. anthracis*.^2,28–36^ Of these few SrtB enzymes studied, not all have been tested for activity *in vitro*, and those that have revealed substantially reduced activity as compared to SrtA proteins from the same organisms.^24,26,30,37^ Overall, substrate preferences for these SrtB enzymes vary, incorporating various residues at all positions: [E/N/P/Q/S/V]-[A/P/V]-[K/Q/S]-[S/T]-[G/N/S]. This is in contrast to the stringent specificity of SrtA enzymes for the LPXTG, or very similar, recognition sequences.^4,38–40^ Notably, work by ourselves and others has discovered additional promiscuity in SrtA enzymes at the P1 and P1’ positions, particularly for SrtA proteins from the *Streptococcus* genera, using *in vitro* experiments.^38–40^ However, identified and putative cellular SrtA substrates are thought to generally follow the LPXTG motif.^38–44^

Sortases contain a conserved structural core. The sortase-fold is defined as an eight-stranded β-barrel.^12,17,45,46^ Flexible loops connect one β-strand to the next and are designated accordingly, e.g., the loop connecting the β6 strand to the β7 strand is the β6-β7 loop. Several of these loops perform an important role in enzyme recognition and specificity, notably the β4-β5, β6-β7, and β7-β8 loops, of which the sequences and lengths can vary widely between sortases of the same class.^13,17,37,41,42,47^ Between classes, there are several regions of additional structural variation.^1,41^ These regions include the N-terminal region before the β-barrel, the β6-β7 loop, the β7-β8 loop, and the C-terminus extending away from the β-barrel. For example, as compared to SrtA enzymes, the SrtB β6-β7 loop is markedly longer and includes an elongated α-helix; this loop is known to contribute to substrate recognition (**Figure 1**).^24,37^ The catalytic domains of SrtB enzymes also contain an extended N-terminal helix, the role of which is not known (**Figure 1**). Finally, the stereochemistry of substrate binding to SrtB remains unclear; although there is a structure of saSrtB bound to the peptidomimetic NPQT* where *= (2*R*,3*S*)-3-amino-4-mercapto-2-butanol, the P4 Asn is making only one contact with saSrtB, limited to a single side chain hydrogen bond with the backbone carbonyl oxygen of T177; this is in contrast to its importance in the recognition motif.^24,37^ Taken together, SrtB enzymes remain an understudied class of the sortase family.

**Figure 1.**
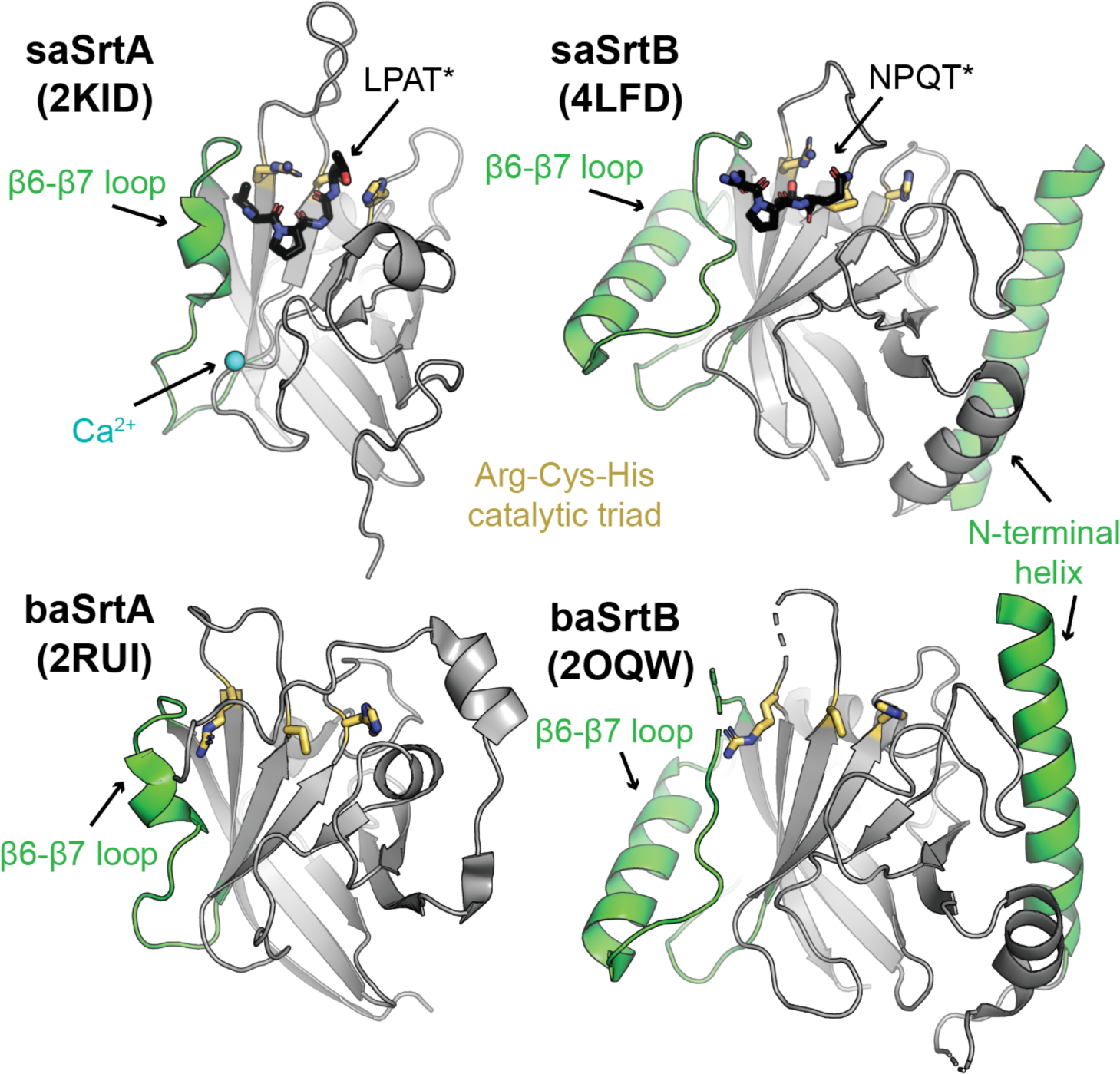
Class A and B sortase structures. The structures of *Staphylococcus aureus* SrtA (PDB ID 2KID) and SrtB (PDB ID 4LFD) enzymes, as well as *Bacillus anthracis* SrtA (PDB ID 2RUI) and SrtB (PDB ID 2OQW) are shown in cartoon representation and colored gray.^24,46,49,74^ Structural features that differ between class A and B sortases are highlighted in green, which include a long N-terminal helix in SrtB and the length of the β6-β7 loop. The side chains of the Arg-Cys-His (left-to-right in this orientation) triad are shown as sticks and colored by atom (N=blue, S=yellow, C=gold). A calcium ion is in blue sphere representation in the saSrtA structure, and peptidomimetics (LPAT* and NPQT*, respectively) for saSrtA and saSrtB are in black sticks and colored by atom (O=red).

To further our understanding of SrtB enzymology, and potentially leverage the activity of these enzymes for protein engineering, in this study we have explored the activity of SrtB enzymes from multiple organisms (*B. anthracis, C. difficile, S. aureus,* and *L. monocytogenes*) along with a series of mutants. Of these, only *B. anthracis* SrtB (baSrtB) revealed reasonable catalytic activity; therefore, we chose to use this enzyme as a model for further characterization. We interrogated the stereochemistry of SrtB substrate-binding using structural modeling with AlphaFold2 and molecular dynamics simulations, as well as mutagenesis. We also tested a variety of nucleophiles for baSrtB. We found that baSrtB requires amino acids outside of the pentapeptide NPKTG binding motif, specifically a P5 Asp and P2’ Asp/P3’ Glu for optimal activity. In addition, a chimeric baSrtB enzyme with the β7-β8 loop sequence from *L. monocytogenes* SrtB revealed an increase of ∼3-fold in relative activity, which we discovered was largely due to a single mutation, A241K. Incorporation of this residue into a A236E/A241K baSrtB double mutant increased activity to 4.4-fold higher than wild-type in a 1.8-hour assay. Finally, we used purified protein substrates to validate sortase-mediated ligation using SrtB enzymes. Overall, our work provides a number of insights into class B sortases and further highlights the potential of these enzymes as intriguing candidates for expanding the catalog of suitable SML substrates and applications.

## Results

### Activity assays of multiple SrtB enzymes

To better understand the relative activities of SrtB enzymes, we first identified a number with available crystal structures in the Protein Data Bank (PDB). These included: *B. anthracis* (baSrtB; PDB IDs 1RZ2, 2OQW),^48,49^ *C. difficile* SrtB (cdSrtB; 4UX7),^32^ *Clostridium perfringens* SrtB (cpSrtB; 5B23, 5YFK),^50^ *L. monocytogenes* (lmSrtB; 5JCV), *S. aureus* SrtB (saSrtB; 1NG5, 1QWZ, 1QX6, 1QXA, 4LFD),^24,25^ and *S. pyogenes* SrtB (spySrtB; 3PSQ).^30^ We reasoned that available monomeric crystal structures, a few of which include either small molecules, a peptidomimetic, or a Gly3 bound, suggested that these proteins would be biochemically tractable to study. However, we chose to exclude additional studies of cpSrtB, as the substrate is not known and was previously shown not to be the heme-transport protein CPE0221,^50^ and spySrtB, due to the additional requirement of the SipA protein cofactor.^51^ Therefore, we moved forward with recombinant expression and purification of the catalytic domains of baSrtB, cdSrtB, lmSrtB, and saSrtB, as described in the Materials and Methods.

With purified proteins in hand, we initially determined relative activities using an established peptide-based assay. Here, putative substrate peptides were synthesized using solid-phase peptide synthesis, as described in the Materials and Methods. Peptide sequences for each protein were as follows, with the pentapeptide recognition motifs underlined: Abz-D<u>NPKTG</u>DEK(Dnp)-*NH_2_* (from *B. anthracis* IsdC (baIsdC)) for baSrtB, Abz-KVE<u>NPQTN</u>AGK(Dnp)-*NH_2_* (from *S. aureus* IsdC (saIsdC)) for saSrtB, Abz-PV<u>PPKTG</u>DSK(Dnp)-*NH_2_* (from CD0386) for cdSrtB, and Abz-T<u>NPKSS</u>DSK(Dnp)-*NH_2_* (from Lmo2186) for lmSrtB, where Abz = 2-aminobenzoyl, Dnp = 2,4-dinitrophenol, K(Dnp) indicates an isopeptide bond with the ε-amine of Lys, and -*NH_2_* indicates a C-terminal amide. When intact, the Dnp moiety quenches fluorescence from the Abz fluorophore via Förster (or fluorescence) resonance energy transfer (FRET); however, upon peptide cleavage by SrtB, the Abz fluorescence is apparent, as described in the Materials and Methods and previously.^16,39,41,42,52,53^ Activity assays revealed that baSrtB was the most active of the four enzymes, while cdSrtB and lmSrtB showed relatively little to no activity (**Figure 2A**). We subsequently tested Abz-K<u>NAKTN</u>DSK(Dnp)-*NH_2_* (from Lmo2185) activity in lmSrtB, which was also a negative result (**Figure S1A**).

**Figure 2.**
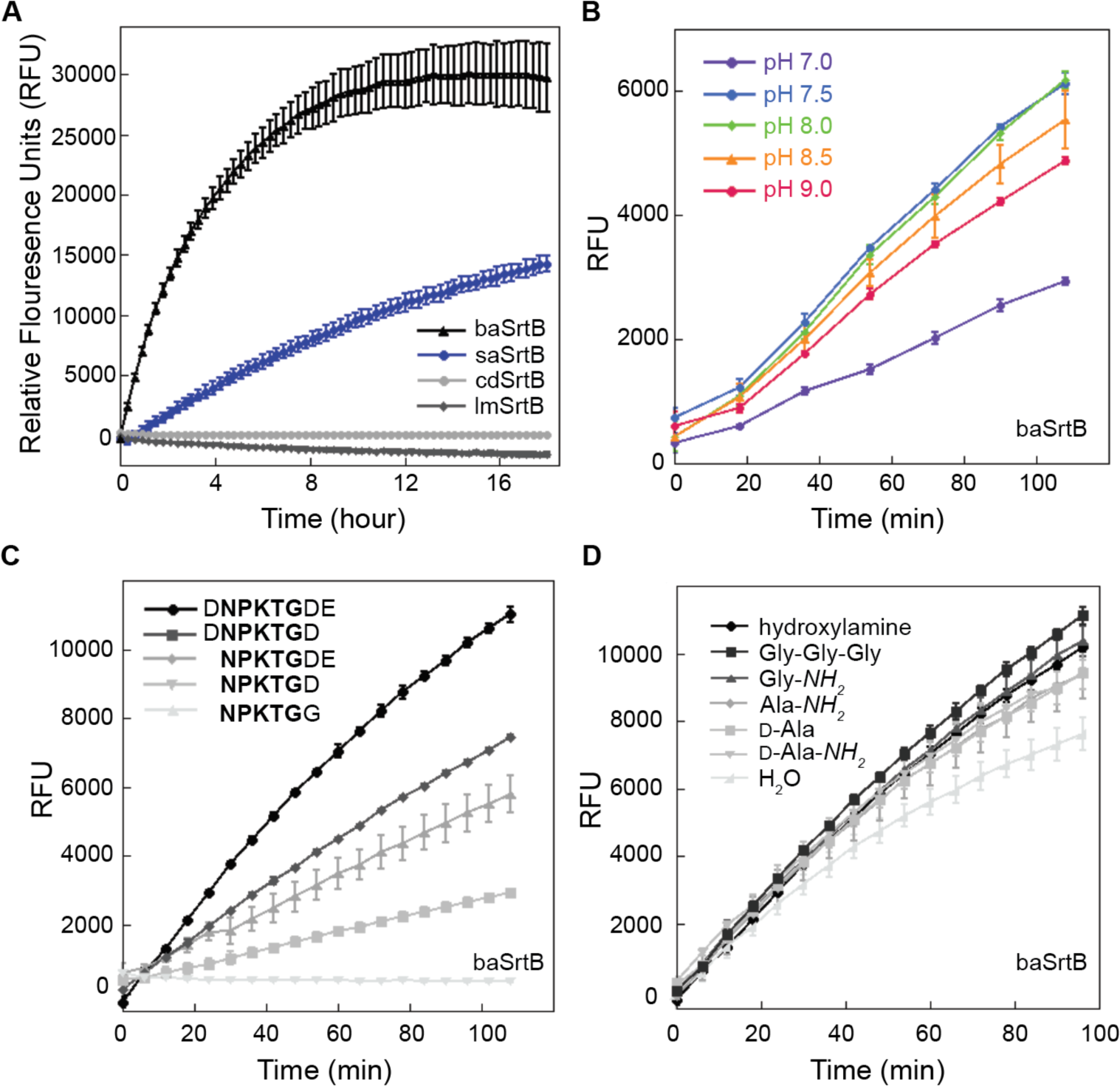
Relative activities of SrtB enzymes. (**A**) A FRET-based reporter assay, with the general structure Abz-(SrtB recognition sequence)-K(Dnp), was used to monitor relative activity of SrtB enzymes from *Bacillus anthracis* (baSrtB), *Staphylococcus aureus* (saSrtB), *Clostridioides difficile* (cdSrtB), and *Listeria monocytogenes* (lmSrtB) organisms. Sequences tested were as follows: Abz-D<u>NPKTG</u>DEK(Dnp)- *NH_2_* (from *B. anthracis* IsdC (baIsdC)) for baSrtB, Abz-KVE<u>NPQTN</u>AGK(Dnp)-*NH_2_* (from *S. aureus* IsdC (saIsdC)) for saSrtB, Abz-PV<u>PPKTG</u>DSK(Dnp)-*NH_2_* (from CD0386) for cdSrtB, and Abz-T<u>NPKSS</u>DSK(Dnp)-*NH_2_* (from Lmo2186) for lmSrtB. (**B-D**) Having established baSrtB as the most active in this assay, additional experiments characterized the pH dependence (**B**), position-specific recognition of substrate (**C**), and preferred nucleophile (**D**) of this enzyme. Wild-type saSrtB and pH-dependence assays were performed in duplicate and all others were run in at least triplicate; all curves represented are averages measured in relative fluorescence units (RFU) following subtraction of negative control (minus SrtB) background fluorescence, and including the standard deviation of replicate experiments.

Our initial assays were run using 50 μM SrtB and 175-200 μM of each respective peptide. This is reflective of the relatively low activities previously observed of SrtB enzymes; in contrast, many of our previous SrtA assays were run with 10 μM enzyme and 50 μM substrate.^41,42^ Because of the long reaction time of these initial SrtB assays as well as the relatively high concentration of the fluorescent peptide, we discovered that we likely encountered *inner filter effects* in our assays, whereby fluorescence is suppressed by a high concentration of fluorophore.^54^ These effects were previously found to affect Abz fluorescence at concentrations as low as 50 μM and we found that the effect become more pronounced over the long timeframe (18 hours) of these initial assays, likely due to evaporation of the reaction mixture.^52^ Therefore, we confirmed our relative activity results for baSrtB, saSrtB, and cdSrtB using an HPLC-based assay over 20-48 h (**Figures S1B-D**), and moving forward, we will restrict our reported FRET-based assay results to 1-2 h, a timeframe which limits these effects.

### Sequence-, pH-, and nucleophile-selectivity of baSrtB

Based on the results of our initial activity assays, we chose to further investigate baSrtB as a model for this class of enzymes. We first wanted to validate previous reports that baSrtB was most active between pH 8.0-9.0, as well as that it recognizes amino acids beyond the NPKTG pentapeptide motif.^26^ We utilized our FRET-based activity assays to test baSrtB with the Abz-D<u>NPKTG</u>DEK(Dnp)-*NH_2_* peptide in the presence of reaction buffers that ranged in pH from 7.0-9.0 in 0.5 pH unit steps (**Figure 2B**). In a 1.8 h assay, our results were similar but in slight contrast to the previously reported results; baSrtB was most active at pH 7.5 and 8.0, with pH 8.5 showing very similar results. We chose to use pH 8.0 for subsequent assays.

To investigate the extended sequence motifs for baSrtB, we evaluated the following peptide sequences (NPKTG pentapeptide is underlined, all contain Abz and K(Dnp-*NH_2_*), which are omitted here for clarity): D<u>NPKTG</u>DE, <u>NPKTG</u>DE, D<u>NPKTG</u>D, <u>NPKTG</u>D, and <u>NPKTG</u>G (**Figure 2C**). Our results confirm that baSrtB requires these three additional residues for full activity. Specifically, we see the largest reductions in activity with the loss of the P5 Asp and P2’ Asp. Notably, the loss of all three residues failed to show any activity in our assay (**Figure 2C**).

Finally, we wanted to test the nucleophile-selectivity of baSrtB, as there is known diversity of the amine nucleophile that serves as the ligation partner for SrtB enzymes. Depending on the organism, this nucleophile may be an N-terminal pentaglycine strand, dialanine, or *meso-*diaminopimelic acid, amongst other possibilities.^55–57^ When a suitable nucleophile is present, then acyl-enzyme formation (the original cleavage reaction) is thought to be the rate-limiting step.^58^ However, in the absence of a suitable nucleophile, the apo-enzyme is restored through hydrolysis. In this case, hydrolysis is expected to become the rate-limiting step and the overall reaction rate is reduced.^58^ We therefore wondered if a poor nucleophile might be a contributing factor to low baSrtB *in vitro* activity. The endogenous nucleophile for baSrtB is thought to be *m*-diaminopimelic acid, though baSrtB may not recognize *m*-diaminopimelic acid *in vitro*.^26^

We tested the following nucleophiles with baSrtB and the substrate Abz-D<u>NPKTG</u>DEK(Dnp)-*NH_2_*: triglycine, Gly-*NH_2_*, Ala-*NH_2_*, D-Ala*-NH_2_*, and D-Ala, as well as the strong nucleophile hydroxylamine (**Figure S2**). For these studies, the D-amino acid nucleophiles D-Ala*-NH_2_* and D-Ala were selected based on structural and chemical similarity to the presumed nucleophilic portion of *m-*diaminopimelic acid, and L-amino Ala-*NH_2_* was selected to investigate any stereochemical preference that might be displayed by the enzyme. Glycine nucleophiles were selected based on previously published experiments demonstrating the ability of baSrtB to catalyze ligation reactions to the nucleophile GGGK-Biotin.^26^ Hydroxylamine was previously shown to serve as a suitable nucleophile for several sortase homologs by ourselves and others; this was used as the nucleophile for the wild-type activity assays (**Figure 2A**).^4,17,38,41^ All reactions progressed at seemingly similar rates in a 1.6 h activity assay regardless of nucleophile, and were only slightly elevated above the activity levels seen in the hydrolysis control (**Figure 2D**). Products formed were confirmed by HPLC-based assays using mass spectrometry (**Table S1**). Taken together, we concluded that our choice of nucleophile was not dramatically affecting the relatively weak baSrtB activity. In addition, our results showed that baSrtB can use L- and D-alanine as a nucleophile, a result not previously tested.

### Stereochemistry of substrate binding to SrtB enzymes

To better understand the results described above and to characterize baSrtB, we wanted to investigate the stereochemistry of substrate binding. We first attempted to use X-ray crystallography to determine a complex structure of catalytically inactive C233A baSrtB protein, expressed and purified as described in the Materials and Methods, bound to a peptide matching the D<u>NPKTG</u>DE sequence. Using previously determined crystallization conditions, as well as positive hits from a number of conditions in commercially available crystallization screens, we were able to generate quality crystals that diffracted with resolution <2 Å (**Figure S3A**). However, data processing, molecular replacement using the apo baSrtB as a model (PDB ID 2OQW), and initial rounds of refinement confirmed that the peptide was not present in our crystal lattice. This was true for crystals grown under multiple salt and precipitant conditions.

We therefore decided to model substrate binding to baSrtB using Alphafold2.^59^ As an input, we used a single sequence matching the full-length *B. anthracis* IsdC, followed by a linker (G_4_S)_2_, then the full-length baSrtB without the initiator Met (see Supporting Information for full sequence used). Although there are no substrate-bound structures of baSrtB in the PDB, we reasoned that the substrate may be situated into the peptide-binding cleft in the predicted folded structure, based on other known sortase structures. Indeed, the output AlphaFold2 models with the 5 highest confidence scores position the NPKTG recognition sequence in the active site of baSrtB (**Figure S3B**). The baIsdC protein contains a long, disordered linker, approximately residue N148 to R213, the residue immediately preceding the transmembrane helix; however, the location of baIsdC in the 5 models does not substantially vary although it is not fixed in one location (**Figure S3B**). Notably, baIsdC is processed by a signal peptidase during membrane translocation, therefore, we also used AlphaFold2 to model this complex using the baIsdC sequence starting at A30. This resulted in an equivalent model for the full-length baSrtB protein (RMSD < 0.2 Å), therefore, we will exclude these residues moving forward in order to better represent the mature folded protein.^27^ Alignment of the main chain atoms of the recognition motif sequence (DNPKTGDE), baIsdC transmembrane helix, and baSrtB revealed RMSD values of < 0.6 Å for all models aligned to the model labeled as ranked_0. We will use ranked_0 as our structural model here due to this high degree of similarity (**Figure 3A**).

**Figure 3.**
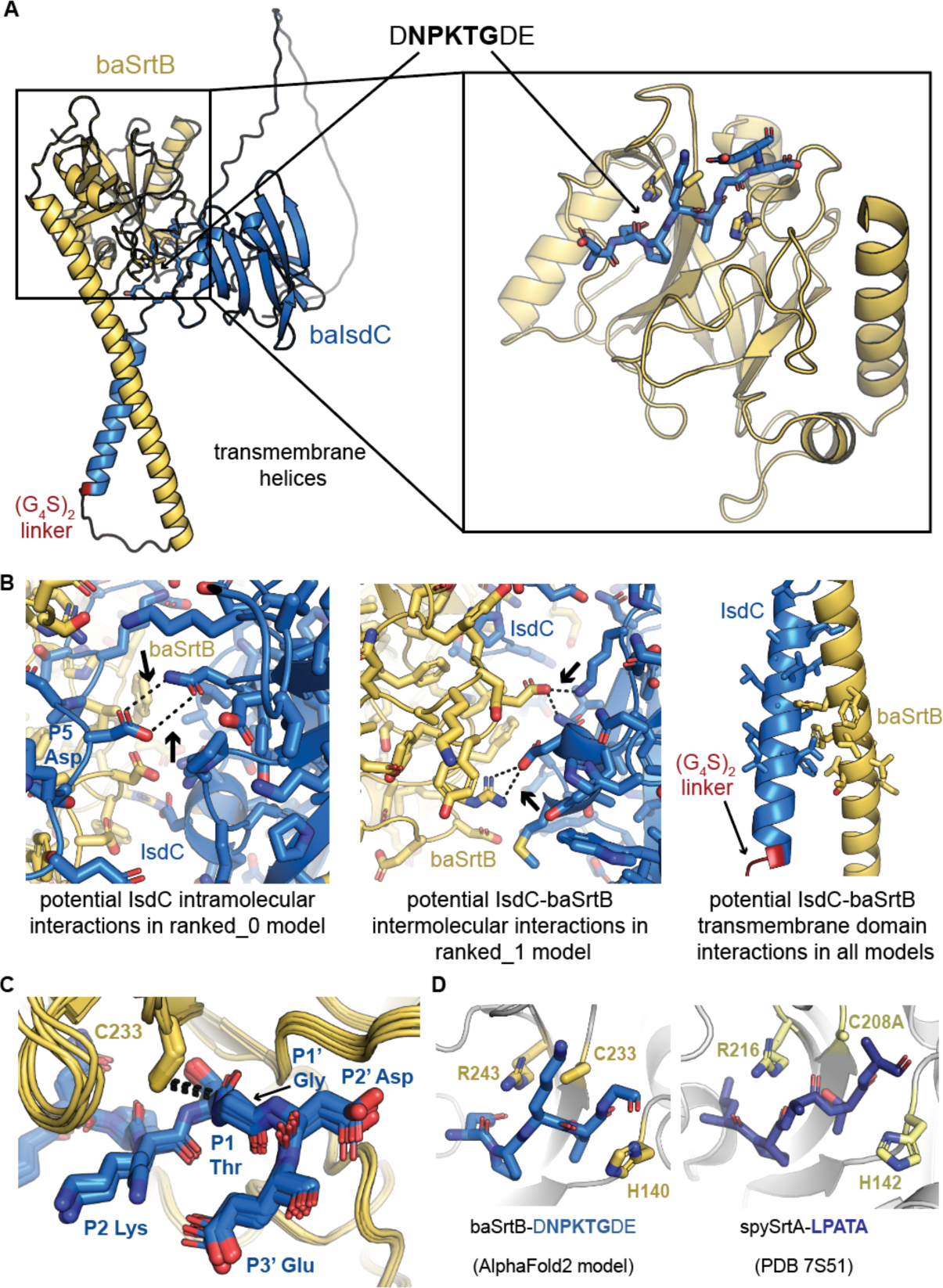
An AlphaFold2 model of baSrtB-substrate binding. (**A**) An AlphaFold2 model of baIsdC binding baSrtB was generated using the input of a single polypeptide sequence containing both full-length proteins, and a (G_4_S)_2_ linker (labeled). The zoomed-in catalytic domain is also shown of baSrtB bound to the DNPKTGDE recognition sequence from baIsdC. (**B**) The 5 AlphaFold2 output models (ranked_0-4) indicated a number of potential interactions which may impact baSrtB activity, as highlighted here and described in the main text. (**C**) All 5 AlphaFold2 models show the recognition sequence from baIsdC in a catalytically-competent conformation with respect to baSrtB. (**D**) BaSrtB-substrate binding is very similar to an experimentally-determined structure of *Streptococcus pyogenes* SrtA (spySrtA) bound to the LPATA peptide (PDB ID 7S51), with respect to orientation of substrate and location of catalytic residues.^17^ For all, protein structures are shown in cartoon representation with side chains modeled as sticks, where applicable, and colored by atom (N=blue, O=red, S=yellow, C=as labeled by protein). BaIsdC, or its substrate sequence, DNPKTGDE, is in blue, baSrtB is in golden yellow, and in (**D**), spySrtA is in yellow-green, and its peptide substrate, LPATA, in purple-blue.

Our AlphaFold2 model of baIsdC-baSrtB allowed us to make a number of predictions about this binding interaction. Although our analyses will be primarily focused on DNPKTGDE binding to baSrtB, we also observed three potential interactions worth further analyses: (1) the folded domain of baIsdC may be forming intramolecular contacts with DNPKTGDE or (2) intermolecular electrostatic contacts with SrtB, and (3) the transmembrane helices may dimerize in the membrane (**Figure 3B**). Any of these interactions could enhance reaction rates and may have a substantial effect on endogenous activity, which has not been measured to our knowledge. With respect to DNPKTGDE recognition by the catalytic domain of baSrtB, the peptide is well-positioned in the active site, in a conformation that we predict is competent for activity (**Figure 3C**). For example, the thiol group of C233 is 3.6 Å from the carbonyl carbon of the P1 Thr in the ranked_0 model, a compatible distance for nucleophilic attack by this rotamer or others. In the other four models, the distances range from 3.5 - 3.8 Å (**Figure 3C**). The other catalytic residues, H140 and R243 are in a consistent position with respect to the peptide as compared to a previously reported spySrtA-LPATA structure (PDB ID 7S51) (**Figure 3D**).^17^

Investigating specific contacts at peptide positions revealed differences compared to a published saSrtB-NPQT* structure (PDB ID 4LFD).^24^ Main chain atoms (641 total) from the A-protomer from saSrtB-NPQT* aligned with an overall RMSD of 0.67 Å (**Figure S4A**). As mentioned above, the P4 Asn in PDB ID 4LFD is forming a single hydrogen bond with the backbone carbonyl oxygen of T177 in saSrtB (**Figure S4B**). In contrast, our model revealed specific interactions with three baSrtB residues, the side chain hydroxyl of Y191 in the β6-β7 loop, the side chain hydroxyl of S231 in the β7 strand, and the side chain guanidino group of the catalytic R243, a characteristic consistent with previous observations by ourselves and others that the P4 residue stabilizes the ligand in SrtA enzymes (**Figure 4A**).^15,17,20^ We also see differences in the position of the P3 Pro; whereas, in saSrtB-NPQT*, this residue is primarily stabilized by I182, in our model of baSrtB-DNPKTGDE, there are hydrophobic interactions observed with side chain atoms of L106, F121, and Y138 (**Figures 4B, S4C**). Notably, all four of these SrtB residues (Ile, Leu, Phe, and Tyr) are structurally conserved in the SrtB sequences we analyzed.

**Figure 4.**
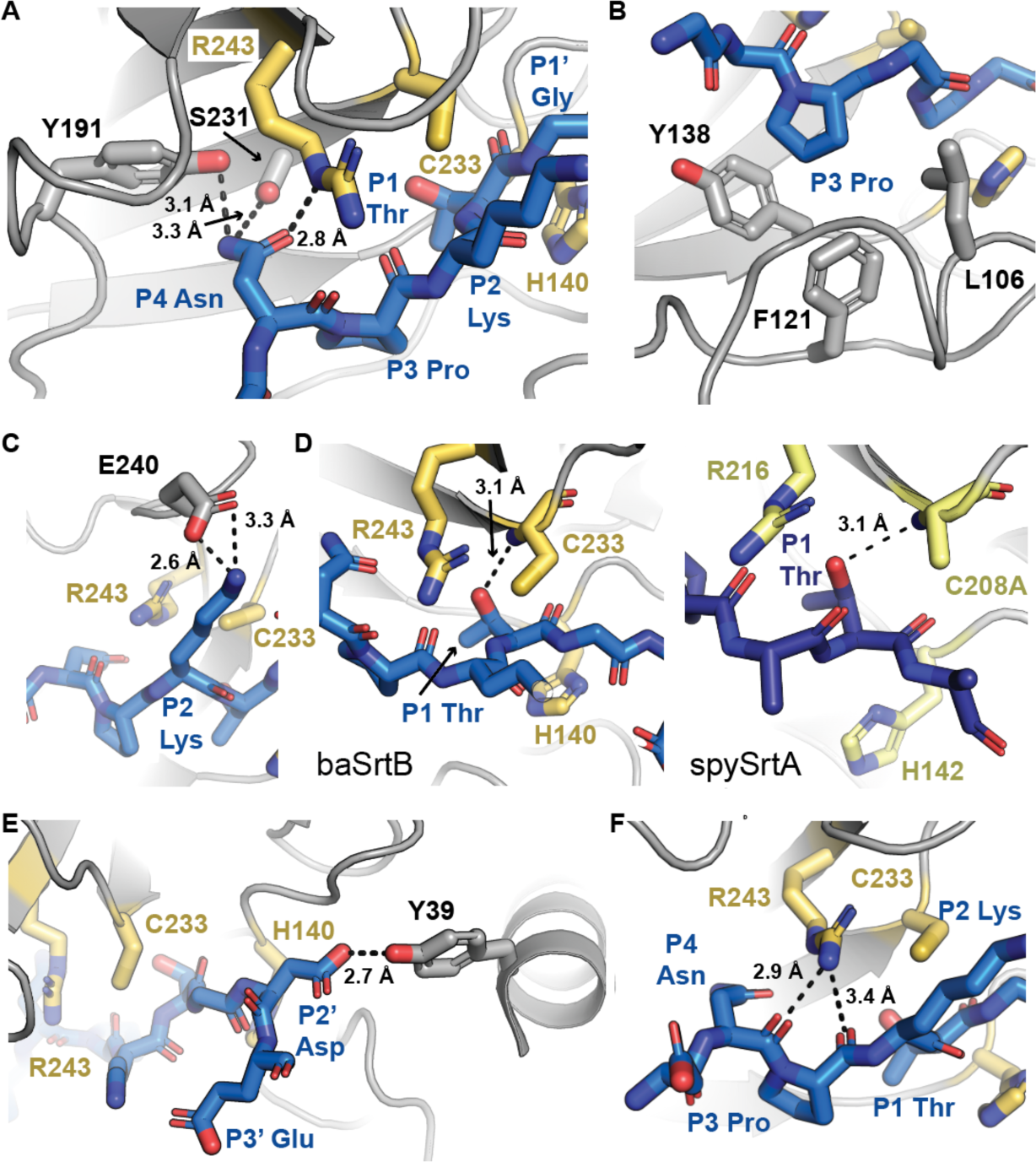
Position-specific recognition of DNPKTGDE peptide substrate by baSrtB. Position-specific recognition of the peptide substrate by baSrtB is highlighted for the (**A**) P4 Asn (D<u>N</u>PKTGDE), (**B**) P3 Pro (DN<u>P</u>KTGDE), (**C**) P2 Lys (DNP<u>K</u>TGDE), (**D**) P1 Thr (DNPK<u>T</u>GDE), and (**E**) P2’ Asp (DNPKTG<u>D</u>E) residues. Backbone interactions between the catalytic Arg and peptide are in (**F**). For all, baSrtB is in gray cartoon, with relevant side chain atoms shown as sticks and colored by atom (N=blue, O=red, S=yellow, C=as labeled by protein). The catalytic residues of baSrtB, H140, C233, and R243, are in golden yellow. The peptide substrate is in blue and colored by atom. In (**D**), spySrtA (PDB 7S51) is in yellow-green, and its peptide substrate, LPATA, in purple-blue for comparison with baSrtB. Measurements are in dashed black lines, with distances labeled.

Additional residues in the binding motif are challenging to compare directly, as the catalytic Cys thiol groups are shifted 6 Å with respect to each other, despite otherwise good alignment in the sortase domains. However, in our baSrtB-DNPKTGDE model, we see electrostatic interactions between the P2 Lys and E240 in the β7-β8 loop (**Figure 4C**), the P1 Thr positioned similarly to that of our spySrtA-LPATA structure and stabilized by the Cys amide (**Figure 4D**), and interactions with the P2’ Asp and side chain atoms of Y39 in the N-terminal helix (unique to SrtB enzymes) and K143 in the β4-β5 loop (**Figure 4E**). We also observed similar stabilizing backbone interactions between the guanidino group of the catalytic R243 residue and the carbonyl oxygens of the P3 Pro and P2 Lys (**Figure 4F**), consistent with the spySrtA-LPATA complex structure.^17^

Interestingly, we did not observe any specific interactions with baSrtB which indicated why the P5 Asp and P3’ Glu in the <u>D</u>NPKTGD<u>E</u> (positions underlined) sequence have an effect on activity (**Figure 2C**). We reasoned that this may be due to the nature of the AlphaFold2 model generated, namely the presence of the full-length baIsdC protein, of which both residues may be at the interface for potential intramolecular interactions in baIsdC. Although we did not observe consistent contacts amongst the AlphaFold2 models between the P5 Asp and folded domain of baIsdC, we wanted to investigate the stereochemistry of these positions further, as well as the stability of the entire recognition motif. Therefore, we ran triplicate molecular dynamics (MD) simulations of the peptide fragment with baSrtB.

Our MD results confirmed that the baSrtB protein and DNPTKTGDE peptide are relatively stable over the course of the three 1000 ns simulations, visualized as the average root mean squared deviation (RMSD) of Cα atoms over time (**Figures 5A, S5A**). For peptide amino acids, there is a clear difference in relative flexibility for the pentapeptide motif, NPKTG, as compared to the P5 Asp and P2’/P3’ D/E residues (**Figure S5B**). While interacting residues for the pentapeptide motif agree with those discussed above, based on the ranked_0 model alone, analyses of electrostatic interactions over the simulations suggests that the P5 Asp most commonly interacts with: N102, Y124, R125, and Y190, the P2’ Asp with: Y39, R116, H140 (catalytic His), K143, and Y235, and the P3’ Glu with: Y39, R116, R141, K143, and K152. There appear to be two dominant positively-charged “pockets” for P2’ and P3’ binding, formed by R116/K143 and R141/K152, respectively. An electrostatic potential surface map of baSrtB confirms that the S2’ and S3’ binding sockets form the most-positively charged surface, consistent with the observed preference for addition of the P2’/P3’ D/E substrate residues (**Figures 2C, 5B**).

**Figure 5.**
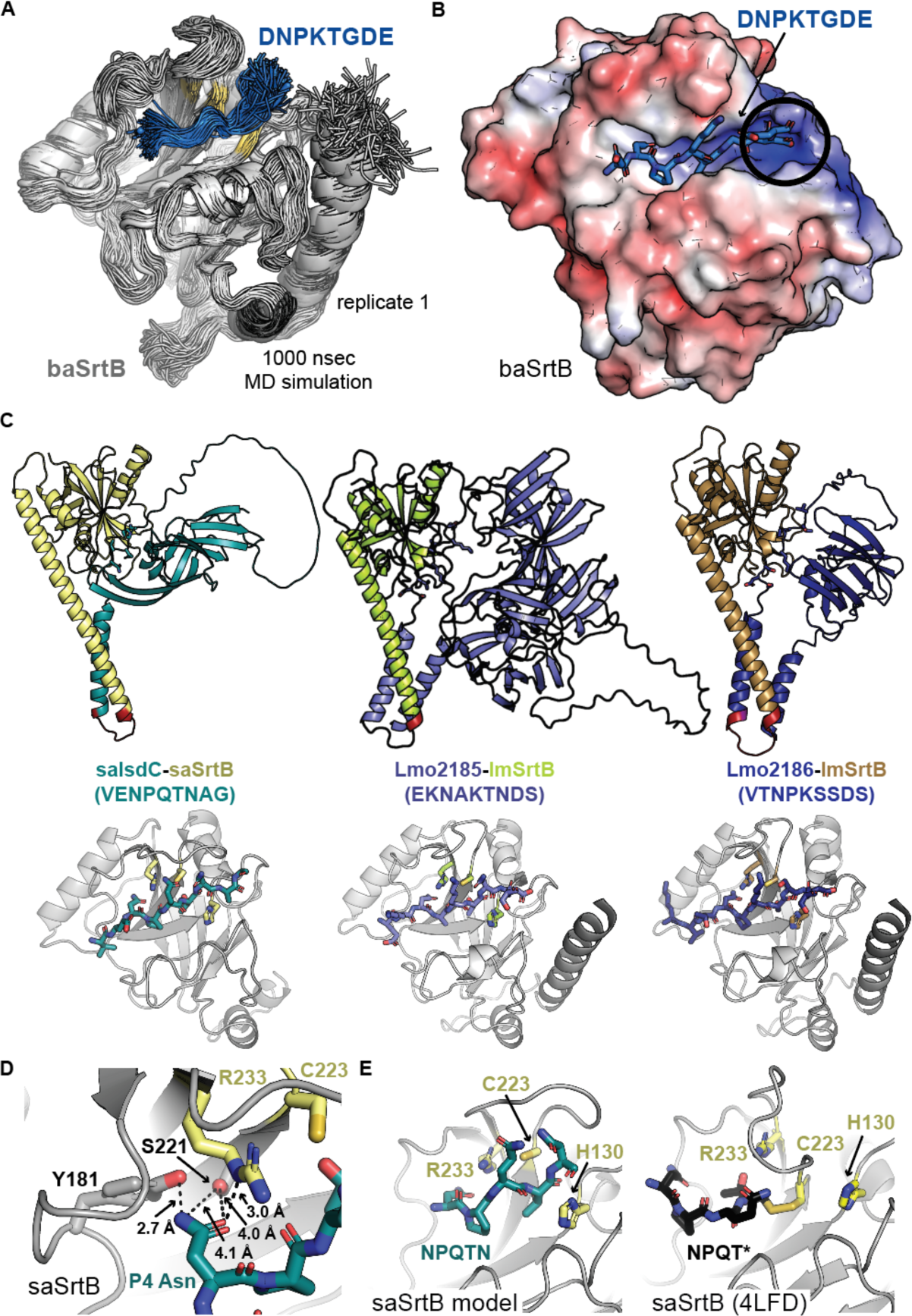
Molecular dynamics simulation of baSrtB-substrate, and AlphaFold2 models of other SrtB enzymes. (**A**) Alignment of 200 states (every 5 nsec of simulation time) during one replicate of our baSrtB-DNPKTGDE molecular dynamics simulation. The baSrtB enzyme is in gray cartoon, with catalytic residues highlighted in golden yellow, and the peptide substrate is in blue cartoon. (**B**) Electrostatic potential surface map (±5 eV, on a red (negative) to blue (positive) scale) of the baSrtB catalytic domain with the DNPKTGDE peptide shown as blue sticks and colored by atom (N=blue, O=red). A positively-charged pocket where we predict the P2’ Asp and P3’ Glu residues interact is circled. (**C**) AlphaFold2 models of saIsdC-saSrtB (left), Lmo2185-lmSrtB (middle), and Lmo2186-lmSrtB (right) are shown in cartoon representation and colored as labeled. The zoomed in catalytic domains bound to peptide substrate (sequences in parentheses) are below each model in cartoon representation, with the catalytic residue side chains and peptide substrates as sticks and colored by atom. (**D**) Predicted interactions between the P4 Asn in the saSrtB substrate, <u>N</u>PQTN, and enzyme. Distances are shown as black dashed lines and labeled. (**E**) Comparison of the saSrtB model generated here (left) and NPQT* peptidomimetic binding in the experimentally-determined structure, PDB ID 4LFD.^24^ Structures are rendered as elsewhere in the figure.

Although relatively inactive in our hands, we also wanted to investigate the stereochemistry of ligand binding to the saSrtB and lmSrtB proteins. Therefore, we created substrate-bound models using AlphaFold2 for saSrtB-VE**NPQTN**AG, lmSrtB-EK**NAKTN**DS, and lmSrtB-VT**NPKSS**DS using a similar protocol as that with baSrtB. Specifically, we modeled single polypeptide sequences of saIsdC-(G_4_S)_2_-saSrtB, Lmo2185-(G_4_S)_2_-lmSrtB, and Lmo2186-(G_4_S)_2_-lmSrtB (**Figure 5C**). The structural models containing saIsdC were treated equivalently as those with baIsdC and again, models of the full-length saSrtB with or without the saIsdC signal sequence included were equivalent (RMSD < 0.1 Å); here, the mature protein is predicted to start at A29.^27^ We also attempted to use this approach for the cdSrtB protein, but the peptide was not properly docked near the active site. All input sequences used in AlphaFold2, including for IsdC-baSrtB, are in the Supporting Information. We also extracted peptide-bound models of these complexes and ran 1000 ns triplicate MD simulations. As with baSrtB, ligand-bound saSrtB and lmSrtB are stable over the course of our trajectories with a well-positioned pentapeptide motif, consistent with the clear non-covalent interactions that stabilize peptide binding in our baSrtB-DNPKTGDE model (**Figures 5C, S5C-E**).

This result was particularly interesting for saSrtB, as there is an available structure bound to a peptide mimetic, NPQT* where T* = (2*R*,3*S*)-3-amino-4-mercapto-2-butanol (PDB ID 4LFD), as mentioned above.^24^ Whereas none of the four protomers of the crystal structure show specific binding of the P4 Asn to saSrtB residues, our model suggests that the P4 Asn may interact with the side chain atoms of Y181 and S221, analogous residues as those predicted to stabilize P4 Asn binding to baSrtB (Y191 and S231, respectively) (**Figure 5D**). Overall, peptide binding is quite different in PDB ID 4LFD versus our AlphaFold2 model (**Figure 5E**). Notably, the NE2 atom of the catalytic His (H130) side chain is 8.9 Å away from the P1 Thr carbon that would be the site of initial nucleophilic attack in the experimental structure (PDB ID 4LFD). Because ligand binding to saSrtB in our model was consistent with that for baSrtB, we wanted to use mutagenesis to further investigate these observations and directly assay the validity of our baSrtB AlphaFold2 and MD results.

### Mutagenesis supports model of ligand binding to baSrtB

To biochemically interrogate our substrate-bound model of baSrtB, we tested 15 mutations and truncations. These included: N-terminal truncations aimed to investigate the role of the additional α-helix in SrtB enzymes, as compared to SrtA (baSrtB_42-254_ and baSrtB_65-254_), N-terminal α-helix point mutations (D38A, Y39A, Y39F), mutations in catalytic residues (H140A, R243A), and several hypothesized to interact with ligand residues (L106A, R116A, Y191A, Y191F, S231A, Y235A, and E240A) (**Figure 6A**). All proteins were recombinantly expressed, purified, and tested as previously described and in the Materials and Methods.

**Figure 6.**
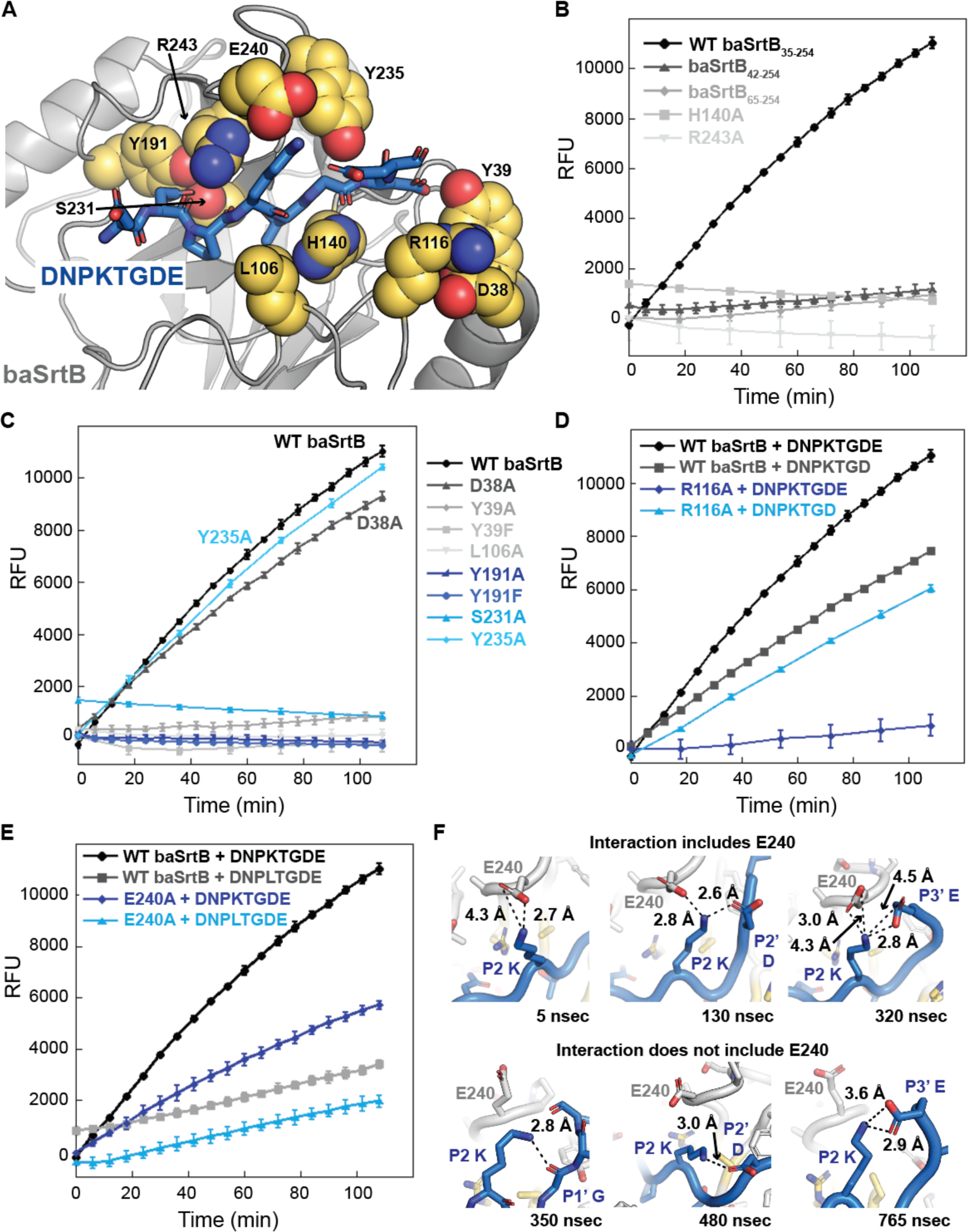
Mutagenesis of baSrtB confirms validity of AlphaFold2 model. (**A**) The baSrtB enzyme is in gray cartoon and DNPKTGDE peptide substrate in blue sticks and colored by atom (N=blue, O=red). Mutations tested in this work are shown as golden yellow spheres, colored by atom, and labeled. (**B-C**) FRET-based activity assays for baSrtB variants are shown as the averages of at least triplicate experiments, with standard deviation. The N-terminal helix truncations and catalytic residue mutations, H140A and R243A, are in (**B**) and all other mutations in (**C-E**). The wild-type data is in both graphs for comparison between the two. (**D**) Comparison of relative activity levels between WT baSrtB and R116A baSrtB with the DNPKTGDE and DNPKTGD peptides, as labeled. Activity is mostly restored in the R116A baSrtB variant when the peptide lacks a P3’ Glu. (**E**) Comparison of relative activity levels between WT baSrtB and E240A with the DNP**K**TGDE and DNP**L**TGDE peptides, as labeled. (**F**) Representative states from one of the replicate MD simulations of WT baSrtB with DNPKTGDE reveals interactions between the peptide P2 Lys and E240 or intrapeptide atoms. Structures are colored as in (**A**) with the peptide main chain shown as cartoon and side chain atoms shown as sticks in all cases but one (bottom left), where the peptide main chain is also in stick representation.

Our mutagenesis data agreed with the baSrtB-DNPKTGDE model. We saw a total loss in activity in the catalytic residue mutations, those predicted to interact with the P4 Asn, Y191 and S231, and L106A, a residue that appears to interact with and stabilize the P3 Pro (**Figures 4B, 6B-C**). In addition, both N-terminal α-helix truncations dramatically reduced activity and point mutations suggested this is predominantly due to interactions with a single residue, Y39 (**Figure 6B-C**). Both Y39A and Y39F baSrtB proteins revealed similar levels of activity, consistent with an interaction between the Tyr side chain hydroxyl group and the P2’ Asp residue that is suggested by our model (**Figures 4E, 6C**). These findings suggest that the extended N-terminal α-helix is important for baSrtB catalysis. The activity of the D38A and Y235A proteins were relatively similar to wild-type (**Figure 6C**). The R116A baSrtB protein revealed a decrease in activity consistent with our structural observations of a positive P2’, P3’ binding pocket, suggesting this may be a primary P3’ Glu-interacting residue (**Figures 5B, 6D**). Consistent with this, the activity of R116A baSrtB with the DNPKTGD peptide, which lacks the P3’ Glu, restored activity to near wild-type levels for this peptide (**Figure 6D**). We did not explicitly test R141, K143, and K152, which we predicted may also contribute to this positive binding pocket for the P2’ Asp and P3’ Glu. In addition, the E240A baSrtB protein showed a loss of activity as compared to wild type baSrtB (**Figure 6E)**. Our modeling suggests the E240 residue forms electrostatic interactions with the P2 Lys (**Figure 4C**). Consistent with this, we see a marked reduction in activity for wild-type baSrtB, and similar levels of activity between wild-type baSrtB and E240A, with the substrate peptide DNP**L**TGDE, which substitutes a Leu for the P2 Lys (**Figure 6E**). However, E240A activity is also reduced with the DNP**L**TGDE substrate compared to DNP**K**TGDE activity, indicating that the P2 Lys may form additional interactions that facilitate activity (**Figure 6E).** Indeed, we visualized several states in one of our representative MD simulations which contained intrapeptide interactions of the P2 Lys with the main chain carbonyl of the P1’ Gly, or side chain atoms of P2’ Asp and P3’ Glu residues (**Figure 6F**). These interactions occurred both in the presence and absence of E240 (**Figure 6F**). Taken together, these results support our substrate-bound baSrtB structure as a good model of the stereochemistry of this complex, specifically for the core NPKTG residues; furthermore, we hypothesize our AlphaFold2 models represent the likely mode of ligand binding for other SrtB enzymes, e.g., saSrtB and lmSrtB.

### Chimeric proteins reveal interactions that dramatically improve baSrtB activity

Previously, we studied the β7-β8 loop of SrtA enzymes in great detail, discovering a number of interactions that affect activity and specificity.^41,42^ We were therefore curious if residues in this loop have a similar effect on SrtB enzymes. To this end, we created a number of chimeric enzymes, with the β7-β8 loop sequences of saSrtB, lmSrtB, and cdSrtB engineered into baSrtB. We recombinantly expressed, purified, and tested these variants as previously described, and as in the Materials and Methods. We will refer to these proteins as: baSrtB_aureus_ (β7-β8 loop sequence: <u>C</u>**ED**A**YSETTK**<u>R</u>), baSrtB_monocytogenes_ (<u>C</u>D**TEK**D**Y**E**K**G<u>R</u>), and baSrtB_difficile_ (<u>C</u>**T**Y**EF**D--**DA**<u>R</u>), where the catalytic Cys and Arg residues are underlined, bold indicates mutations as compared to baSrtB, and dashes indicate changes in loop length. For reference, the baSrtB β7-β8 loop sequence is: <u>C</u>DYALDPEAG<u>R</u>.

All three chimeric enzymes were active, although to varying degrees (**Figure 7A**). Whereas the baSrtB_aureus_ and baSrtB_dificile_ chimeras revealed reduced activity as compared to wild-type baSrtB, we saw a dramatic increase with baSrtB_monocytogenes_; here, the t = 1.8 h fluorescence was 2.8-fold higher in the β7-β8 loop variant as compared to wild-type baSrtB (**Figure 7A**). We made single point mutations to interrogate this result further, changing one baSrtB β7-β8 loop residue (sequence: D<u>YAL</u>D<u>P</u>E<u>A</u>G) at a time to the corresponding amino acid in the lmSrtB loop (sequence: D<u>TEK</u>D<u>Y</u>E<u>K</u>G, differences underlined): Y235T, A236E, L237K, P239Y, and A241K (note: we are using baSrtB numbering because of the consistency in loop lengths between these enzymes) (**Figure 7B**). All proteins were recombinantly expressed, purified, and tested as previously reported and as in the Materials and Methods. Of these mutant proteins, P239Y baSrtB and Y235T baSrtB had relatively equivalent activity, and A236E, L237K, and A241K baSrtB revealed increased activity over wild-type baSrtB (**Figure 7B**). Specifically, the increase in activity of A241K baSrtB recapitulated, and even surpassed (3.6-fold over wild-type at t = 1.8 h), that seen in baSrtB_monocytogenes_ (**Figure 7B**). A double mutant, A236E/A241K baSrtB, increased this effect further, with a 4.4-fold increase in activity as compared to wild-type at t = 1.8 h (**Figure 7B**).

**Figure 7.**
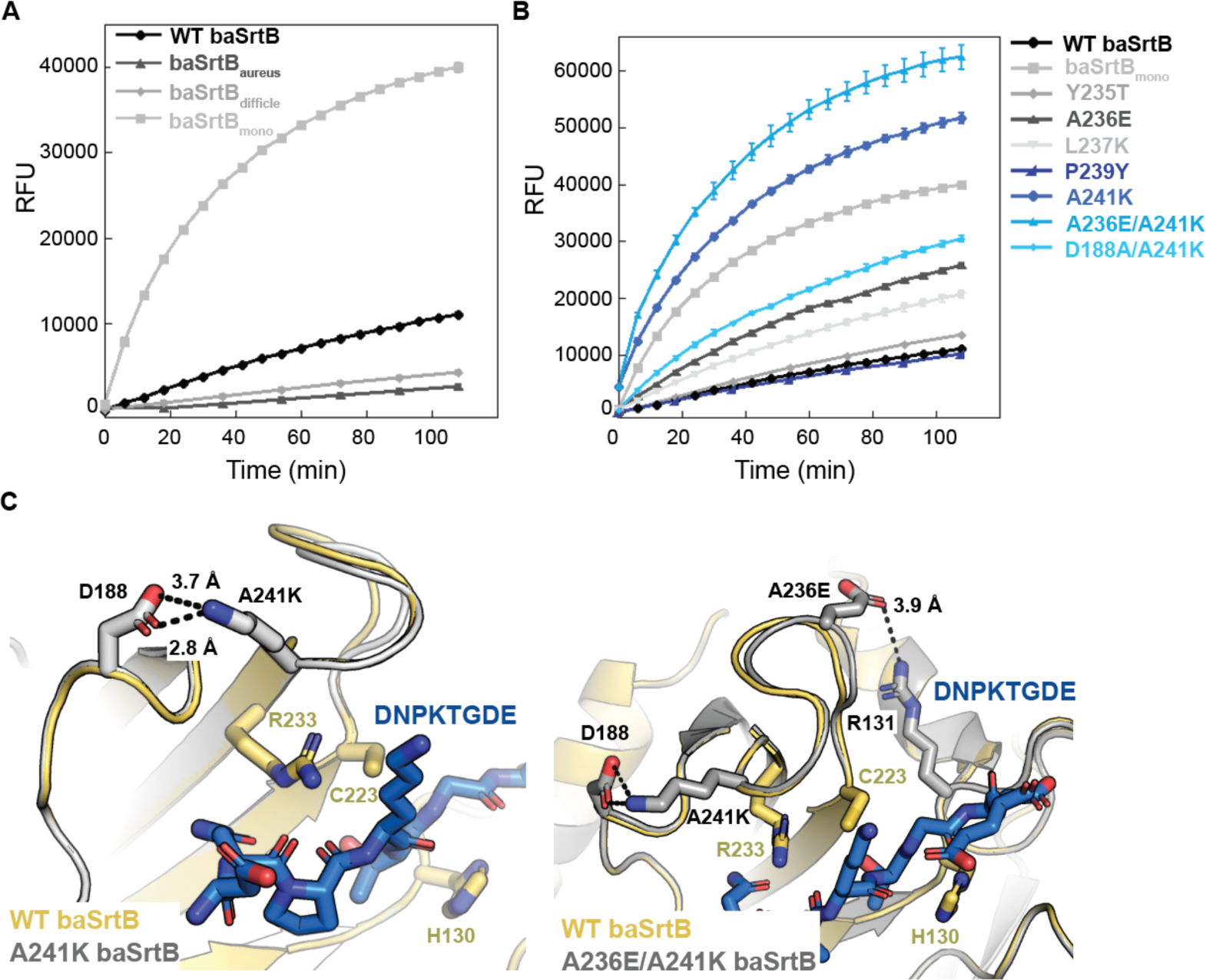
Loop swapped chimeras elucidate baSrtB mutants with higher relative activities. (**A**) BaSrtB chimeras were created by inserting the β7-β8 loop sequences from saSrtB (baSrtB_aureus_), cdSrtB (baSrtB_difficle_), and lmSrtB (baSrtB_mono_), and FRET-based activity assays were run in at least triplicate. The averaged values are shown, along with standard deviation. (**B**) Single mutations were created in baSrtB to replicate amino acids in the β7-β8 loop of lmSrtB. The results of averaged replicate experiments are shown, with standard deviation. The wild-type baSrtB and baSrtB_mono_ data is also shown for comparison. Double mutants were also tested to assay additive effects (A236E/A241K) or to investigate the basis of the activity difference (D188A/A241K). (**C**) Hypothesized interactions that increase the relative activity of A241K and A236E/A241K baSrtB as compared to wild-type are shown using AlphaFold2 models created of the variant proteins. The wild-type baSrtB protein is in golden cartoon, with the A241K (left) or A236E/A241K (right) baSrtB proteins in gray cartoon, as labeled. Relevant side chain sticks are shown as gray sticks and colored by atom (N=blue, O=red). The DNPKTGDE peptide substrate is in blue sticks and colored by atom. Measurements are shown as black dashed lines, and distances labeled.

AlphaFold2 models generated of baIsdC-(G_4_S)_2_-A241K and baIsdC-(G_4_S)_2_-A236EA241K baSrtB indicated that the observed effect may be due to stabilizing interactions between A236E with R131 and/or A241K with D188, respectively (**Figure 7C**). While the β7-β8 loop backbone atoms of the three proteins (wild-type, A241K, and A236E/A241K) are very similar, we predict that these stabilizing interactions may affect conformational dynamics during catalysis. We expressed, purified, and tested the double mutant D188A/A241K baSrtB, which revealed ∼2-fold decreased activity as compared to A241K baSrtB alone, although it is still 2.1-fold higher than wild-type (**Figure 7B**). Taken together, our results revealed single (A241K) and double (A236E/A241K) mutations that have a large effect on baSrtB activity.

### Ligation experiments with baSrtB

Finally, we wanted to test the ability of baSrtB to ligate two proteins together. We initially designed two fluorescent substrate proteins, mTurq2-DNPKTGDEGGGG and G(G_4_S)_2_-SYFP2. Expression and purification details are in the Materials and Methods. As described in the Materials and Methods, we ran a ligation assay using similar concentrations to those from our peptide-based experiments described above. We saw the clear appearance of a ligation product using SDS-PAGE and mass spectrometry (**Figures 8A-B, S6, Table S2**). We ran this experiment in triplicate using the wild-type, A241K, and A236E/A241K baSrtB enzymes.

**Figure 8.**
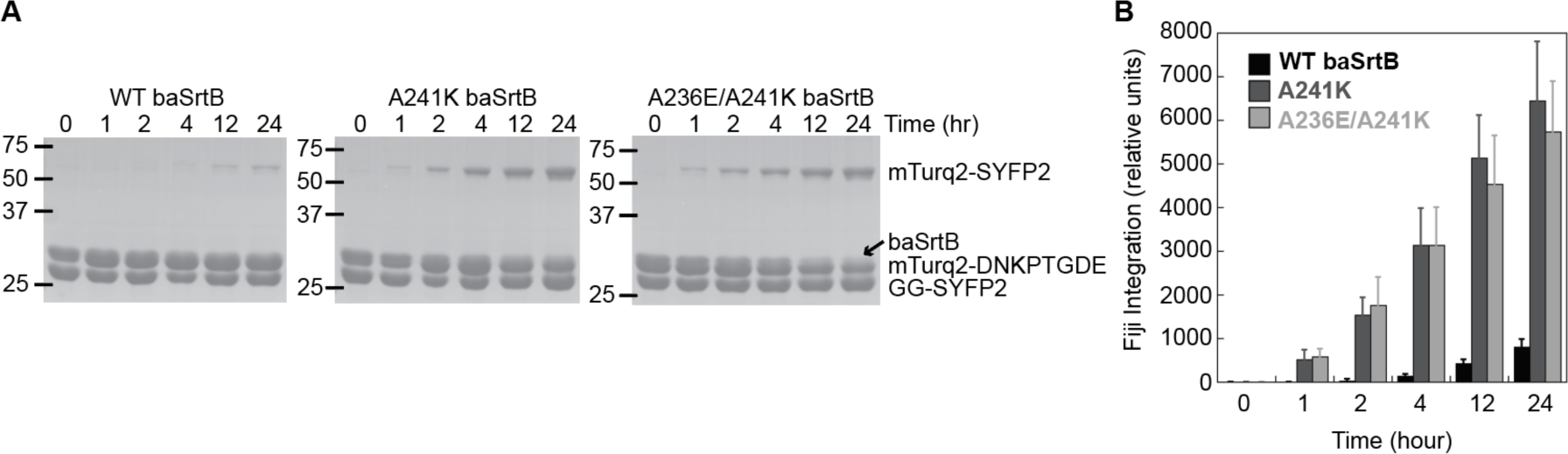
Ligation assays using purified components show baSrtB sortase-mediated ligation. (**A**) SDS-PAGE gels revealed that 50 μM wild-type, A241K, and A236E/A241K baSrtB enzymes are capable of ligating 200 μM mTurq2-DNPKTGDEGGGG and 200 μM GG-SYFP2 proteins together. All protein species are labeled to the right of the A236E/A241K gel. One replicate is shown, and gels from two additional replicate experiments are in the Supporting Information. (**B**) Fiji (<u>F</u>iji is <u>J</u>ust ImageJ) was used to quantify protein bands from all three experiments as relative integration units. Background normalization was not done due to the limitation of separating the baSrtB and mTurq2 protein bands, as described in the main text, leading to higher error values than expected from visualization of the gels alone; however, the increased ligation activities of A241K and A236E/A241K baSrtB as compared to wild-type are clear based on this quantification, which is also consistent with our FRET-based peptide activity assays.

Quantification of our ligation products was challenging due to the inability to use the baSrtB protein band as a normalizing control. The baSrtB protein is at approximately the same molecular weight as mTurq2, and we were unable to differentiate between the two for quantification. Despite this, we see clear mTurq2-SYFP2 ligation mediated by SrtB, the first time to our knowledge that a class B sortase has been used in an SML reaction to link two protein domains. Furthermore, non-background normalized quantification based solely on the mTurq2-SYFP2 protein band agreed with the gels visualized by eye, as well as our previous assays, in that the A241K and A236E/A241K baSrtB proteins are more active than wild-type (**Figures 7B, 8A**). Here, ligation product appears earlier (t = 1 h) and there is ultimately more produced over the 24 h assay (**Figure 8B**). Taken together, we provide evidence that baSrtB is indeed a viable enzyme for use in SML applications, and that we have identified a single mutation (A241K) that can greatly enhance the efficiency of its use for protein engineering purposes.

## Discussion

Despite 25 years of research into bacterial sortases, class A sortases remain the best studied of this enzyme family. Of this work, the vast majority investigates the characteristics, mechanism, and SML utilization of *Staphylococcus aureus* SrtA, which arguably contains some unique traits, e.g., dependence on allosteric activation by calcium and strict specificity for the LPXTG binding motif.^1,4,38–40,46^ While *S. aureus* SrtA has relatively high activity with respect to other studied sortases, previous directed evolution experiments revealed that a small number of mutations, e.g., a pentamutant, can have dramatic effects on overall catalytic efficiency, increasing it by >100-fold.^60^ In addition, biochemical and structural studies by ourselves in recent years investigated the structurally-conserved β7-β8 loop of sortases, specifically amongst *Streptococcus* SrtA proteins, finding that residues in this loop can have a major impact on the specificity and activity of a given enzyme.^13,17,41,42,53^ Here, we chose to turn our attention to the understudied class B sortases, which had previously been shown to have relatively poor activity as compared to class A sortases from the same organisms.^24,26,30,37^ Based on initial experiments with 4 enzymes, we chose *Bacillus anthracis* SrtB to use as a model enzyme to investigate general properties of this sortase class and to test if our previous approach of engineering β7-β8 loop chimeras would elucidate a better understanding of class B sortases.

Our results revealed a number of new insights into baSrtB and class B sortases in general. As described here, baSrtB recognizes amino acids beyond the pentapeptide motif, here NPKTG, typically used to describe sortase recognition. Specifically, a P2’ Asp is required for baSrtB activity, while P3’ Glu and P5 Asp residues also dramatically contribute (**Figure 2C**). Furthermore, a single mutation in the N-terminal helix, a structural feature that differentiates class B and class A sortase catalytic domains, Y39, can dramatically reduce activity confirming it is necessary for full baSrtB activity; we hypothesize this is due to interactions with the P2’ residue (**Figures 4E, 6B-C**). Due to a lack of general activity, we were unable to test if there are similar non-covalent interactions formed in the other SrtB enzymes studied and their respective P2’ target residues.

We also used AlphaFold2 to model the substrate-sortase interaction for several SrtB enzymes by utilizing a single polypeptide sequence combining the full-length primary sequences of substrate and SrtB with a Gly-Ser linker. Although additional tools to predict the structures of complexes which utilize AlphaFold have been introduced in recent years, e.g., ColabFold or AlphaFold – Multimer, this approach was able to take advantage of freely available AlphaFold2 servers (here, the European Galaxy Server) and quickly predict the sortase-substrate structure.^59,61–63^ Furthermore, our models suggested possible interactions between the transmembrane domains of both proteins, which we are currently testing. We hypothesize that this additional site of substrate recognition could increase reaction rates substantially in bacteria, and may prove useful for SML applications.

Our models largely agree with the binding conformation of available SrtA-peptide structures, and seem to be catalytically competent based on what is known about the general sortase reaction mechanism.^1,21^ While the only available structure of a class B sortase, *S. aureus* SrtB, bound to a peptidomimetic, NPQT*, raised questions about the lack of specific contacts with the P4 Asn in the binding motif, our models for 3 SrtB enzymes suggested that this amino acid is stabilized by non-covalent bonds with conserved Tyr (β6-β7 loop) and Ser (β7 strand) residues, as well as the catalytic Arg (**Figure 4A**). We used mutagenesis to verify that knocking out any one of these residues abrogates binding for baSrtB and a D<u>N</u>PKTGDE peptide (Asn is underlined) (**Figures 6B-C**). These models also support earlier studies by ourselves and others that the catalytic Arg likely stabilizes binding of the peptide substrate as opposed to direct contact with tetrahedral oxyanion intermediates, as previously proposed.^15,17,46^ Instead, we argue that a highly conserved Thr residue, immediately preceding the catalytic Cys, as well as backbone amides, stabilize the acyl-enzyme intermediate, which is consistent with the structural models presented here as well.^17^

Finally, following identification of more active baSrtB variants via our β7-β8 loop chimera experiments, we tested the potential of SrtB for SML. We successfully showed that baSrtB can facilitate ligation of two proteins *in vitro* (**Figure 8**). We identified single (A241K) and double (A236E/A241K) baSrtB variants with 4-fold increased activity over a ∼2-hour period in our activity assays, as well as a several-fold increase in a sortase-mediated ligation experiment, using this approach. This suggests that a more extensive mutagenesis screen, e.g., using directed evolution as was previously done with SrtA, could uncover SrtB variants with even better catalytic efficiencies relatively easily. Considering the varied recognition sequences between SrtB enzymes, between each other and as compared to class A sortases, this may prove incredibly useful for SML applications. Taken together, our work provides additional insight into class B sortases and acts as a proof of principle that understudied sortases, or classes of sortases, may hold great potential for the next generation of sortase-based protein engineering tools.

## Materials and Methods

### Expression and purification of recombinant proteins

Accession numbers of SrtB catalytic domains used for expression and purification include: saSrtB_30-244_ (UniProt SRTB_STAA8), baSrtB_35-254_ (UniProt A0A6L8PZR0_BACAN), cdSrtB_26-255_ (NCBI WP_167414662.1), lmSrtB_26-246_ (UniProt SRTB_LISMO). All baSrtB variants were based on this wild-type sequence. Proteins were expressed and purified largely as previously described for SrtA.^17,41,42^ Briefly, His-tagged sequences, which also contained a TEV protease cleavage site, were inserted into pET28a(+) plasmids (Genscript) and transformed into BL21 (DE3) *Escherichia coli* cells. Following growth and overexpression of the protein of interest using IPTG, cells were resuspended in lysis buffer [0.05 M Tris pH 7.5-8.0, 0.15 M NaCl, 0.5 mM ethylenediaminetetraacetic acid (EDTA)] and lysed by sonication. Purified protein was isolated by immobilized metal affinity chromatography (IMAC) using a 5 mL HisTrap HP column (Cytiva) with wash buffer [0.05 M Tris pH 7.5-8.0, 0.15 M NaCl, 0.02 M imidazole pH 7.5-8.0, and 0.001 M tris(2-carboxyethyl)phosphine (TCEP)] and elution buffer [wash buffer with 0.3 M imidazole pH 7.5-8.0]. Eluted protein was characterized by SDS-PAGE, and pure fractions were concentrated using Amicon Ultra 10K Centrifugal Filters. Final purification was achieved by size exclusion chromatography (SEC) with running buffer [0.05 M Tris pH 7.5, 0.15 M NaCl, 0.001 M TCEP for initial assays in **Figures 2A-B, S1B-D** or 0.1 M Tris pH 8.0, 0.3 M NaCl]. Protein concentrations were determined using theoretical extinction coefficients calculated using ExPASy ProtParam.^64^ Purified proteins were flash-frozen using liquid nitrogen and stored at -80 °C.

The His tag of baSrtB protein used for crystallography was cleaved off using TEV protease. BaSrtB and TEV protease were incubated at a 1:100 ratio at 4 °C overnight. This was followed by a second IMAC purification step, with the buffers described above, prior to SEC. Crystallization attempts consisted of 1-1.5 mM C233A baSrtB protein in a 1:1 ratio with *Ac*-DNPKTGDE-*NH_2_* peptide. Following incubation and crystallization by hanging drop vapor diffusion using a mixture of 2 μL protein + 2 μL well solution, crystals appeared in a few days to weeks under several conditions. Examples included, (1) 0.225 M potassium acetate, 18% (*w/v*) PEG 3350; (2) 0.175 M sodium acetate, 20% (*w/v*) PEG 3350; (3) 0.21 sodium chloride, 22% (*w/v*) PEG 3350, 0.1 M Hepes pH 7; (4) 0.175 M sodium bromide, 22% (*w/v*) PEG 3350. However, in all cases, cryo X-ray diffraction data revealed that the peptide was not bound, as described in the main text.

The mTurq2-DNPKTGDE and GG-SYFP2 sequences were also inserted into the pET28a(+) plasmid (Genscript) and included N-terminal His_6_ and TEV cleavage site purification tags. Expression and purification of mTurq-DNPKTGDE and GG-SYFP2 followed a similar protocol as that described above, with the exception that the SEC running buffer for mTurq-DNPKTGDE was 0.05 M Tris pH 8.0, 0.15 M NaCl, 0.001 M TCEP and for GG-SYFP2, all buffers were at pH 7.5, including the SEC running buffer [0.05 M Tris pH 7.5, 0.15 M NaCl, 0.001 M TCEP].

### Peptide synthesis and FRET-based sortase activity assays

All substrate peptides were synthesized using manual Fmoc solid phase peptide synthesis (SPPS) and contained N-terminal 2-aminobenzoyl (Abz) and C-terminal 2,4-dinitrophenyl lysine (K(Dnp)) moieties, as previously described.^17,38,41,42^ Activity assay reactions were performed in black, flat-bottom Costar 96-well plates. Activity assay reactions were prepared with 0.05 mM SrtB, unless noted below, 0.175-0.2 mM peptide model substrate, unless noted below, 5 mM nucleophile (e.g., hydroxylamine or the others tested), 0.05 M Tris-HCl pH 7.0-9.0 (baSrtB = pH 7.5 then pH 8.0 based on pH experiment; saSrtB = pH 8.5; cdSrtB = 7.5; lmSrtB = pH 8.0), and 0.15 M NaCl. For the data in **Figures S1B-D**, 0.01 mM baSrtB and saSrtB and 0.02 mM cdSrtB were used, with 0.05 mM peptide model substrate. For the data in **Figure 2B**, 0.03 mM baSrtB was used, with 0.055 mM *Abz*- DNPKTGDEK(Dnp)-*NH_2_*. Peptide stocks often contained DMSO to aid solubility. Residual DMSO in reaction mixtures was ≤1.5%, which was confirmed to not affect baSrtB activity. Reactions were monitored by measuring fluorescence intensity (*λ*_ex_ = 320 nm, *λ*_em_ = 420 nm) using a Biotek Synergy H1 plate reader. All reactions were performed at least in triplicate, unless indicated otherwise. Background-subtracted fluorescence (measured in relative fluorescence units, RFU) was obtained by subtracting the RFU of negative controls lacking SrtB. Data was plotted using KaleidaGraph, v5.01 (Synergy Software).

### Ligation assays and analysis using SDS-PAGE

The ligation assays were performed using purified protein components at the following final concentrations: 50 μM baSrtB for all variants used, 200 μM mTurq-DNPKTGDE, and 200 μM GG-SYFP2. Running buffer [0.1 M Tris pH 8.0, 0.3 M NaCl] was added to reach a final buffer concentration of 0.05 M Tris pH 8.0, 0.15 M NaCl. Ligation reactions were incubated at room temperature for 24 h, with timepoints taken from a single reaction solution. The ligation reaction was quenched using SDS sample buffer and subjected to analysis using SDS-PAGE.

### Molecular dynamics simulations

Sortase catalytic domains, equivalent to those expressed and purified for *in vitro* experiments, and the substrate residues indicated as the binding substrate were extracted from determined AlphaFold2 models. The peptide substrate was reidentified as a separate chain. Molecular dynamics simulations were performed in triplicate using GROMACS 2022.4 with the AMBER99SB-ILDN protein, nucleic AMBER94 force fields.^65–69^ PyMOL was used to cap the peptide and protein N-terminus with an acyl group, while the protein C-terminus was capped with N-methyl. The system was solvated with TIP3P water molecules and centered within a cubic box with periodic boundary conditions and a minimum of 1 nm from between the protein and the edge of the box. Ions were added to a 0.15 M physiological ion concentration with a neutral net charge balanced with sodium cations and chloride ions. The steepest descent energy minimization was performed on the solvated system with a maximum force tolerance of 1000 kJ/mol/nm for all structures. Long-range electrostatic interactions were treated with the particle mesh Ewald (PME) algorithm using a grid spacing of 0.16 nm and a 1.0 nm cutoff for Coulombic and Lennard-Jones interactions.^70^ All systems and replicates were equilibrated separately in an NVT ensemble for 100 picoseconds with an integration time step of 2 fs at a reference temperature of 300K with position restraints on all protein heavy atoms with velocities assigned randomly from a Maxwell-Boltzmann distribution. We used the stochastic velocity rescaling thermostat. All systems and replicates were equilibrated separately in an NPT ensemble for 5 nanoseconds without position restraints at a reference pressure of 1.0 bar using the Isotropic Parrinello-Rahman barostat.^71^ All bonds to hydrogen atoms were constrained using the LINCS algorithm.^72^ Sufficient equilibration was verified by a converged protein RMSD and system density. All systems and replicates were simulated in a separate production run in an NVT ensemble for 1000 nanoseconds at a reference temperature of 300K. Molecular dynamics simulations were viewed using PyMOL (Schrödinger software).

### Characterization of activity and ligation assays using liquid chromatography mass spectrometry (LC-MS)

BaSrtB, saSrtB, and cdSrtB peptide activity assays were monitored over 20-48 hours through RP-HPLC using a Dionex Ultimate 3000 HPLC system and a Phenomenex Kinetex 2.6 μM C18 100 Å column (100 x 2.1 mm) [aqueous (95% water, 5% MeCN, 0.1% formic acid) / MeCN (0.1% formic acid) mobile phase at 0.3 mL/min, method: hold 10% MeCN 0.0-1.0 min, linear gradient of 10-90% MeCN 1.0-7.0 min, hold 90% MeCN 7.0-9.0 min, linear gradient of 90-10% MeCN 9.0-9.1 min, re-equilibrate at 10% MeCN 9.1-13.4 min)]. The HPLC system was interfaced with an Advion CMS expression^L^ mass spectrometer in order to confirm substrate and product identity through electrospray ionization mass spectroscopy (ESI-MS). For LC-MS characterization of baSrtB nucleophile activity assay products, reaction solutions were collected from the 96-well plate after approximately 20 hours of incubation time. Protein was removed using Amicon Ultra 3K Centrifugal Filters and flow through was analyzed by LC-MS. For these assays, separation was achieved using a Dionex Ultimate 3000 HPLC system and a Phenomenex Kinetex 2.6 μM Polar C18 100 Å column (100 x 2.1 mm) [aqueous (95% water, 5% MeCN, 0.1% formic acid) / MeCN (0.1% formic acid) mobile phase at 0.3 mL/min, method: hold 0% MeCN 0.0-1.0 min, linear gradient of 0-90% MeCN 1.0-7.0 min, hold 90% MeCN 7.0-9.0 min, linear gradient of 90-0% MeCN 9.0-9.1 min, re-equilibrate at 0% MeCN 9.1-13.4 min)]. As noted above, the HPLC system was interfaced with an Advion CMS expression^L^ mass spectrometer for identification of reaction products by electrospray ionization mass spectroscopy (ESI-MS). LC-MS characterization of sortase-mediated ligation assays was performed using an Agilent 6545XT AdvanceBio Q-TOF system interfaced with an Agilent 1290 HPLC system. Separations upstream of the Q-TOF were achieved with a Phenomenex Aeris^TM^ 3.6 mM WIDEPORE C4 200 Å column (100 x 2.1 mm) [H_2_O (0.1% formic acid) / MeCN (0.1% formic acid) mobile phase at 0.3 mL/min, method: hold 10% MeCN 0.0-1.0 min, linear gradient of 10-90% MeCN 1.0-9.0 min, hold 90% MeCN 9.0-11.0 min, linear gradient of 90-10% MeCN 11.0-11.1 min, re-equilibrate at 10% MeCN 11.1-15.0 min]. Deconvolution of protein charge ladders was achieved using Agilent MassHunter BioConfirm software (version 10.0).

### Structural modeling using AlphaFold2 and software used for structural analyses

Substrate sequences used for modeling from UniProt include: saIsdC (ISDC_STAA8), baIsdC (UniProt A0A2A7DBZ7_BACAN), Lmo2185 (HBP2_LISMO), and Lmo2186 (HBP1_LISMO). The accession numbers of the full-length SrtB sequences are the same as above. Structural models were determined with AlphaFold2 on the European Galaxy server, using default settings.^59,63,73^ Input sequences are in the Supporting Information. The output structures are ranked according to the predicted long-distance difference test (pLDDT).^59^ Output structures were analyzed using PyMOL (Schrödinger software), and unless otherwise noted, the ranked_0 output was used as the reference structure. The electrostatic potential surface map of baSrtB was determined using the APBS plugin in PyMOL.

## Author Contributions (CRediT Classification)

**Sophie N. Jackson:** Conceptualization (equal), Formal Analysis (lead), Investigation (lead), Visualization (supporting), Validation (supporting), Writing – Original Draft Preparation (supporting). **Jadon M. Blount:** Investigation (supporting), Formal Analysis (supporting), Funding Acquisition (supporting), Visualization (supporting), Validation (supporting), Writing – Review & Editing (supporting). **Kayla A. Croney:** Investigation (supporting), Formal Analysis (supporting), Visualization (supporting), Validation (supporting), Writing – Review & Editing (supporting). **Darren E. Lee:** Investigation (supporting), Formal Analysis (supporting), Visualization (supporting), Validation (supporting), Writing – Review & Editing (supporting). **Justin W. Ibershof:** Investigation (supporting), Formal Analysis (supporting). **Kyle Whitham:** Investigation (supporting). **James McCarty:** Formal Analysis (supporting), Funding Acquisition (supporting), Resources (supporting), Writing – Review & Editing (equal). **John M. Antos:** Conceptualization (equal), Visualization (equal), Resources (equal), Supervision (supporting), Writing – Review & Editing (supporting). **Jeanine F. Amacher**: Conceptualization (equal), Funding Acquisition (lead), Visualization (equal), Resources (lead), Supervision (lead), Writing – Original Draft Preparation (lead).

## Acknowledgements

We want to thank all additional members of the Amacher and Antos labs, as well as Dr. Sierra Cullati, for helpful discussion and research support. We also gratefully acknowledge the support of Sarina Kiesser in running mass spectrometry assays. The QTOF system was acquired via an NSF MRI grant (DBI-1920340) to J.M. Antos. This work was additionally supported by NSF CHE-2044958 and a Cottrell Scholar Award from the Research Corporation for Science Advancement to J.F. Amacher, and NSF CHE-2102189 to J. McCarty. J.M. Blount was supported by a Cottrell Postbac Award. Although we were unable to determine a peptide-bound structure of baSrtB bound to a target sequence, we want to thank the Berkeley Center for Structural Biology (BCSB) for crystallography resources. The BCSB is supported in part by the National Institutes of Health, National Institute of General Medical Sciences, and the Howard Hughes Medical Institute. The Advanced Light Source is supported by the Director, Office of Science, Office of Basic Energy Sciences, of the U.S. Department of Energy under Contract No. DE-AC02-05CH11231. The Pilatus detector was funded under NIH grant S10OD021832.

