## Supplemental Material for "Biochemical characterization of *Bacillus anthracis* sortase B: Use in sortase-mediated ligation and substrate recognition dependent on residues beyond the canonical pentapeptide binding motif for sortase enzymes"

Table of Contents:

**Table S1.** LC-MS analysis of N-terminal products from baSrtB nucleophile experiment. 2

**Table S2.** LC-MS analysis of ligation assay components. 3

**Figure S1.** LmSrtB activity assay with alternative substrate sequence and LC-MS analysis of SrtB assays. 4

**Figure S2.** Potential baSrtB nucleophiles. 5

**Figure S3.** X-ray diffraction of baSrtB crystals and AlphaFold2 structural models. 6

**Figure S4.** Comparison of baSrtB model with experimental saSrtB-NPQT* experimental structure (PDB 4LFD). 7

**Figure S5.** Molecular dynamics simulations of SrtB AlphaFold2 models. 8

**Figure S6.** SDS-PAGE gels of baSrtB ligation assays (replicates 2 and 3). 9

**Sequences used for AlphaFold2 modeling.** 10

**Table S1. LC-MS analysis of N-terminal products from baSrtB nucleophile experiment.**

| Nucleophile | Product | Calculated (*m/z*) | Observed (*m/z*) |
|---|---|---|---|
| $H_2O$ | Abz-DNPKT***-OH*** | 693.31 | *not observed* |
| $NH_2OH$ | Abz-DNPKT***-NHOH*** | 708.32 | 708.4 |
| Gly-Gly-Gly | Abz-DNPKT**GGG** | 864.38 | 864.5 |
| Gly-*$NH_2$* | Abz-DNPKT**G-*$NH_2$*** | 749.35 | 749.4 |
| Ala-*$NH_2$* | Abz-DNPKT**A-*$NH_2$*** | 763.37 | 763.5 |
| D-Ala-*$NH_2$* | Abz-DNPKT**A-*$NH_2$*** | 763.37 | 763.4 |
| D-Ala | Abz-DNPKT**A** | 764.35 | 764.4 |

Calculated and observed masses represent $[M+H]^+$ ions (monoisotopic). [-*$NH_2$* = C-terminal primary amide].

**Table S2. LC-MS analysis of ligation assay components.**

| Protein | Calculated (Da) | Observed (Da) |
|---|---|---|
| **Enzymes** | | |
| WT baSrtB | 28159.40 | 28159.71 |
| A241K baSrtB | 28216.50 | 28216.87 |
| A236E/A241K baSrtB | 28274.73 | 28275.55 |
| **Substrates** | | |
| mTurq2-DNPKTGDEGGGG | 28872.25 | 28870.59 |
| $G(G_4S)_2$-SYFP2 | 27721.14 | 27719.33 |
| **Ligation Product** | | |
| mTurq2-SYFP2 | 56045.91 | 56043.76 |

Calculated masses represent average MW and observed masses were determined from deconvolution of the corresponding protein charge ladder.

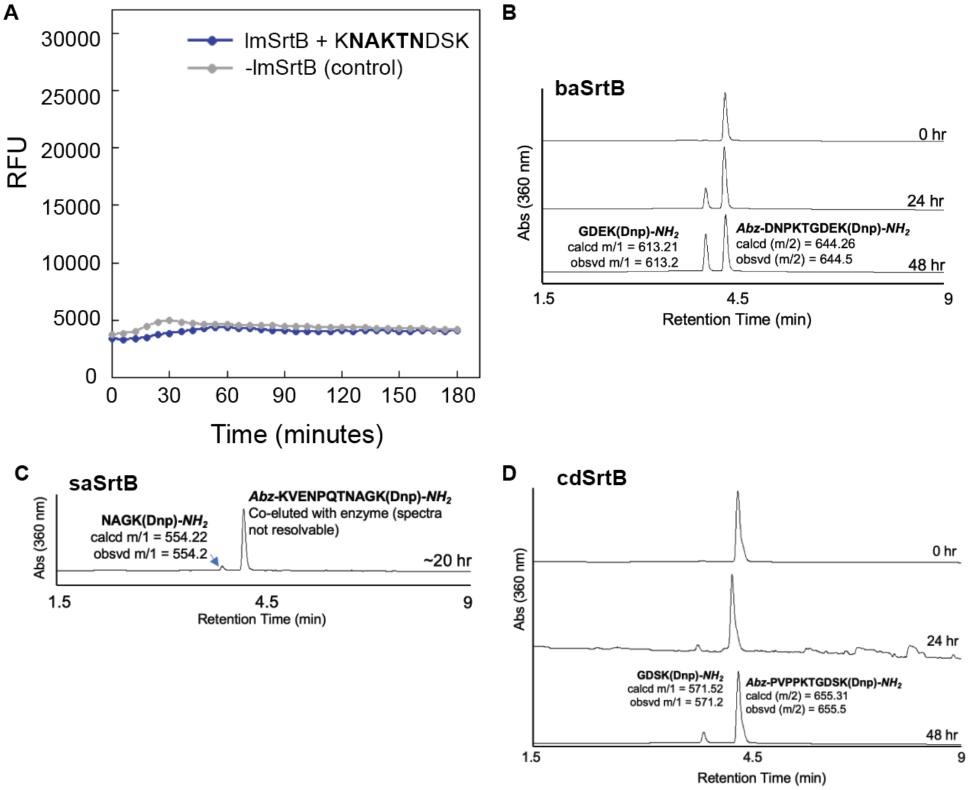

**Figure S1. LmSrtB activity assay with alternative substrate sequence and LC-MS analysis of SrtB assays.** (**A**) Averaged activity assays, in triplicate, of lmSrtB and KNAKTNDSK substrate sequence (from Lmo2185). The negative control (minus lmSrtB) is shown as well. (**B-D**) Analyses using liquid chromatography and mass spectrometry (LC-MS) of (**B**) baSrtB, (**C**) saSrtB, and (**D**) cdSrtB activity assays using peptide substrates. Peaks are labeled.

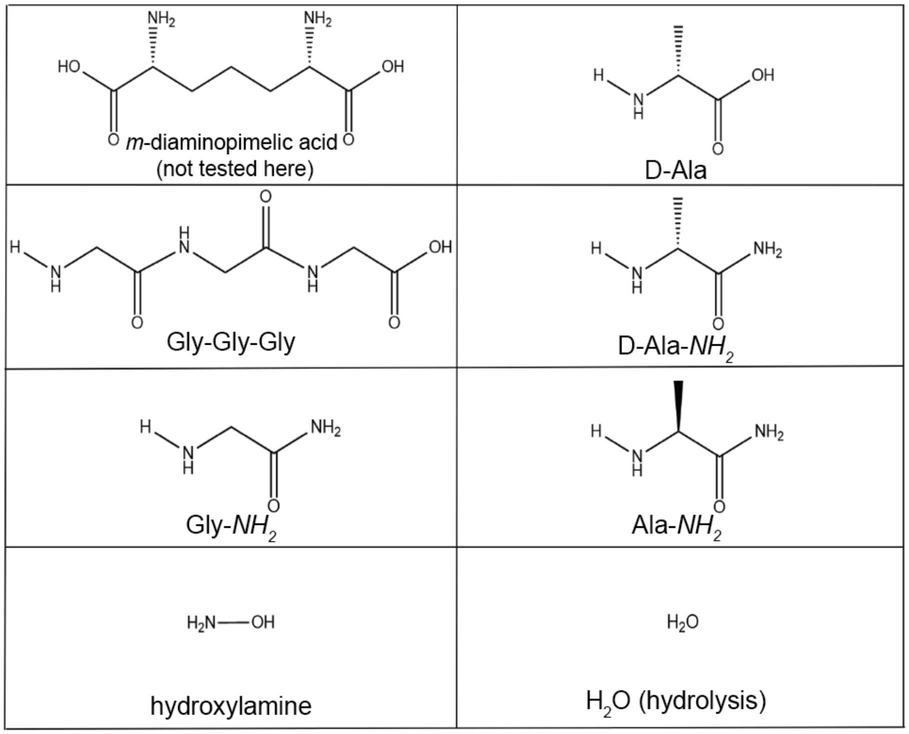

**Figure S2. Potential baSrtB nucleophiles.**

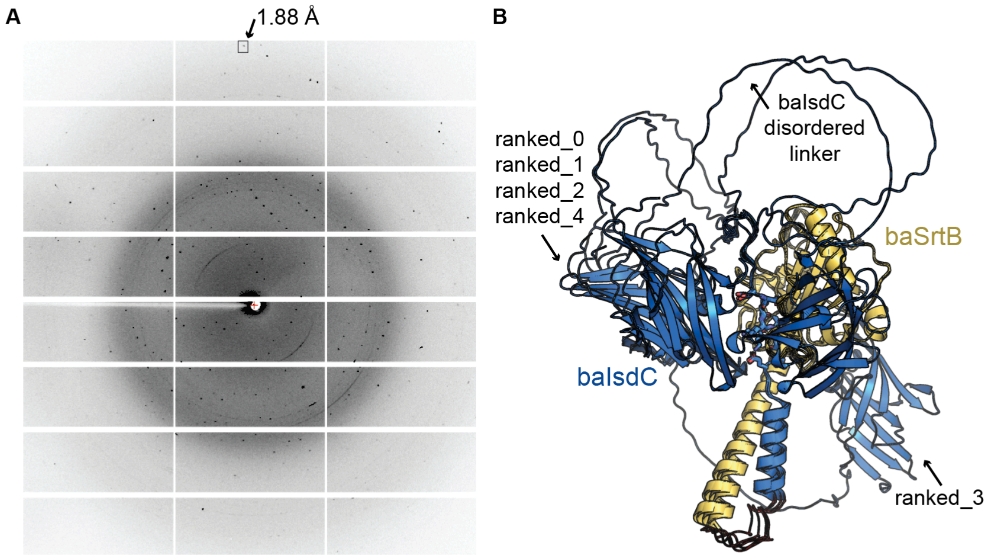

**Figure S3. X-ray diffraction of baSrtB crystals and AlphaFold2 structural models.** (**A**) X-ray diffraction pattern of baSrtB-DNPKTGDE crystals. Subsequent data processing and structure solution revealed that the peptide was not present in the crystals. (**B**) All 5 AlphaFold2 output models are shown for baIsdC-($G_4S$)$_2$-baSrtB. Models are aligned using the full-length baSrtB sequence. An identified baIsdC disordered linker is labeled.

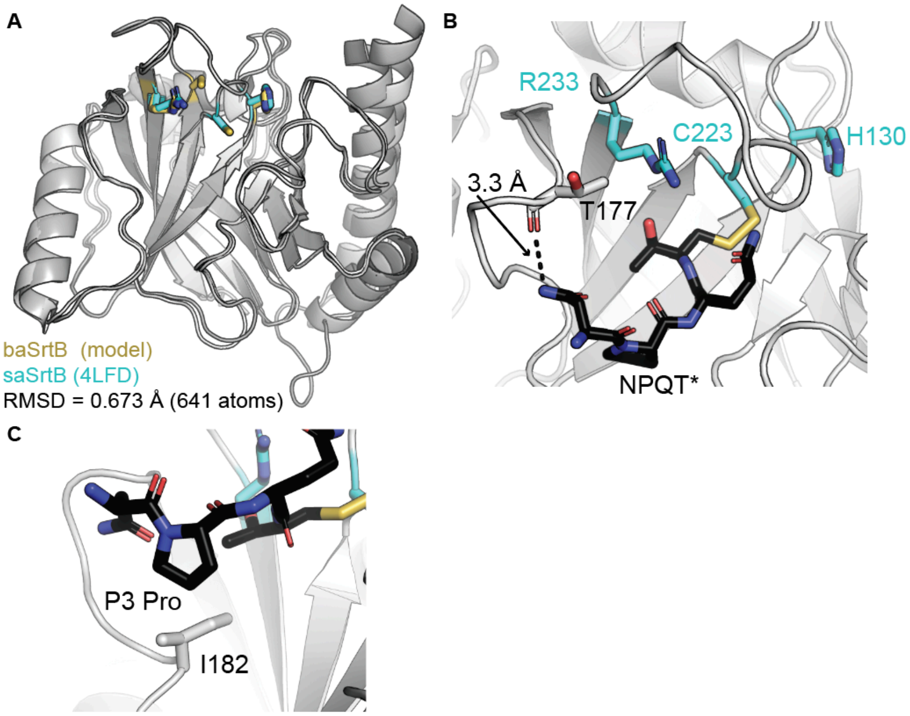

**Figure S4. Comparison of baSrtB model with experimental saSrtB-NPQT* experimental structure (PDB 4LFD).** For all, SrtB proteins are shown in gray cartoon representation, with the side chains of the catalytic triad shown as sticks and colored golden yellow (baSrtB) or cyan (saSrtB) and by atom (N=blue, S=yellow). The NPQT* peptidomimetic is shown as black sticks and colored by atom (O=red). (**A**) Alignment of the main chain atoms of the catalytic domains of baSrtB and saSrtB revealed an overall RMSD = 0.673 Å (641 atoms). (**B-C**) Specific interactions of NPQT* and saSrtB in the experimental structure are highlighted for the (**B**) P4 Asn and (**C**) P3 Pro residues.

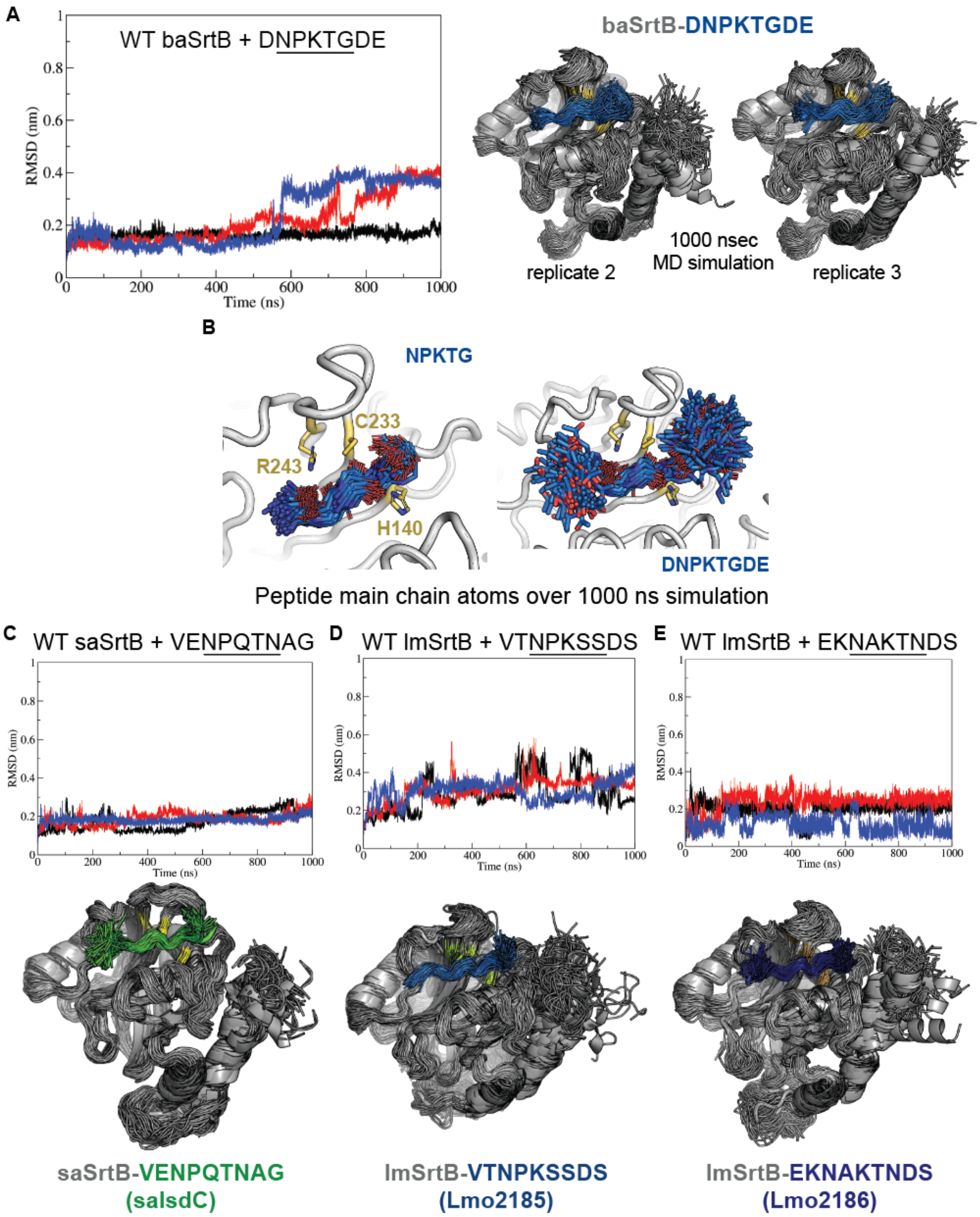

**Figure S5. Molecular dynamics simulations of SrtB AlphaFold2 models.** (**A, C-E**) Average RMSD values for the SrtB C$_\alpha$ atoms over the 1000 nsec trajectories are shown for (**A**) baSrtB, (**C**) saSrtB, (**D-E**) lmSrtB with different substrates. All 3 replicates are graphed. For baSrtB, 200 frames (every 5 nsec) of the remaining 2 replicates are aligned and shown as cartoons and colored as in **Figure 5A**. For saSrtB and lmSrtB, 200 frames (every 5 nsec) from only one replicate for each MD simulation is aligned and shown, with the peptide substrate colored as labeled. (**B**) The peptide backbone atoms are shown as sticks and colored by atom (N=blue, O=red). The peptides of 200 frames (every 5 sec) are aligned to highlight the stability of the pentapeptide motif, NPKTG (left), as compared to the P5 Asp, P2' Asp, and P3' Glu residues (DNPKTGDE, right). The side chain atoms of the catalytic triad residues are shown in golden yellow, colored by atom (S=yellow), and labeled.

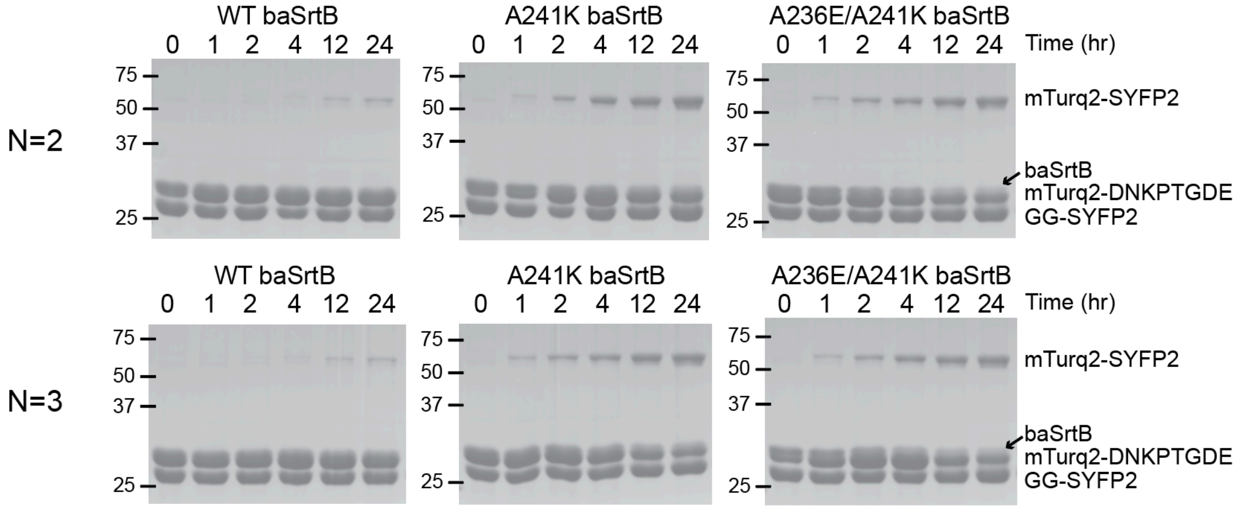

**Figure S6. SDS-PAGE gels of baSrtB ligation assays (replicates 2 and 3).** SDS-PAGE gels from replicate experiments of the baSrtB ligation assay are shown and labeled as in **Figure 7**. These gels were used in the quantification shown in **Figure 8B**.

**Sequences used for AlphaFold2 modeling.** All sequences are in the format: substrate-**$(G_4S)_2$**-SrtB. Accession numbers used for each sequence are in the Materials and Methods. Membrane localization sequences that are subsequently cleaved by a signal peptidase for saIsdC and baIsdC are in italics.

>saIsdC-$(G_4S)_2$-saSrtB
*MKNILKVFNTTILALIIIIATFSNSANA*ADSGTLNYEVYKYNTNDTSIANDYFNKPAKYIKKNGKLYVQITVNHSHW
ITGMSIEGHKENIISKNTAKDERTSEFEVSKLNGKIDGKIDVYIDEKVNGKPFKYDHHYNITYKFNGPTDVAGANAP
GKDDKNSASGSDKGSDGTTTGQSESNSSNKDKVENPQTNAGTPAYIYAIPVASLALLIAITLFV**GGGGSGGGGS**FLT
IVQILLVVIIIIFGYKIVQTYIEDKQERANYEKLQQKFQMLMSKHQEHVRPQFESLEKINKDIVGWIKLSGTSLNYP
VLQGKTNHDYLNLDFEREHRRKGSIFMDFRNELKNLNHNTILYGHHVGDNTMFDVLEDYLKQSFYEKHKIIEFDNKY
GKYQLQVFSAYKTTTKDNYIRTDFENDQDYQQFLDETKRKSVINSDVNVTVKDRIMTLSTCEDAYSETTKRIVVVAK
IIKVS

>baIsdC-$(G_4S)_2$-baSrtB
*MRKISVLPAFIITFVCMLAFLVMPYGKVS*AQLADGTYDINYVIQKAENDSASMANDYFEKPAKLVVKNGEMRVQIPM
NHSAWITEFKAPENGNFVDAKVVNKDESADKRTVEFKIDDLSKPAAAKIHVVVPNVNYDHNYTIRFAFDANVKAVGG
ENKATAVTKNNDQTKTDTKVKEEVKKEESKEVNKEANKGTNESGKAEKTDNPKTGDEARIGLFAALILISGVFLI**GG**
**GGSGGGGS**SSEKERKKKIFFQRILTVVFLGTFFYSVYELGDIFMDYYENRKVMAEAQNIYEKSPMEEQSQDGEVRKQ
FKALQQINQEIVGWITMDDTQINYPIVQAKDNDYYLFRNYKGEDMRAGSIFMDYRNDVKSQNRNTILYGHRMKDGSM
FGSLKKMLDEEFFMSHRKLYYDTLFEGYDLEVFSVYTTTTDFYYIETDFSSDTEYTSFLEKIQEKSLYKTDTTVTAG
DQIVTLSTCDYALDPEAGRLVVHAKLVKRQ

>Lmo2186-$(G_4S)_2$-lmSrtB
MKKVLVFAAFIVLFSFSFLSTGLTAQAALKDGTYSVDYTVIQGDSDSASMANDYFDKPATVTVNGGKSTVSLQVNHS
KWITGLWVEGNAVSVTSKNASSDTRKVSFPVSTLSNPVNAKIKVDIDDDDLNYHHEYQIKLRFDEGSAKALAGAVKS
SDNNTTTPATKSDSSNKVTNPKSSDSSQMFLYGIIFVATGAGLILL**GGGGSGGGGS**LTLVVLGVFLFSGWKIGMELY
ENKHNQTILDDAKAVYTKDAATTNVNGEVRDELRDLQKLNKDMVGWLTIIDTEIDYPILQSKDNDYYLHHNYKNEKA
RAGSIFKDYRNTNEFLDKNTIIYGHNMKDGSMFADLRKYLDKDFLVAHPTFSYESGLTNYEVEIFAVYETTTDFYYI
ETEFPETTDFEDYLQKVKQQSVYTSNVKVSGKDRIITLSTCDTEKDYEKGRMVIQGKLVTK

>Lmo2185-$(G_4S)_2$-lmSrtB
MKKLWKKGLVAFLALTLIFQLIPGFASAADSRLKDGGEYQVQVNFYKDNTGKTTKESSEADKYIDHTATIKVENGQP
YMYLTITNSTWWQTMAVSKNGTRPEKPAQADVYQDRYEDVQTVSTDAAKDTRVEKFKLSSLDDVIFSYMHIKVDAIS
YDHWYQVDLTIDPSTFKVISEPAVTTPVTLSDGIYTIPFVAKKANDDSNSSMQNYFNNPAWLKVKNGKKMVAMTVND
NKTVTALKTTLAGTLQDVKVVSEDKDANTRIVEFEVEDLNQPLAAHVNYEAPFNGSVYKGQADFRYVFDTAKATAAS
SYPGSDETPPVVNPGETNPPVTKPDPGTTNPPVTTPPTTPSKPAVVDPKNLLNNHTYSIDFDVFKDGTTETSMMESY
VMKPALIKVENNQPYVYLTLTNSSWIKTFQYKVNGVWKDMEVVSGDINKNTRTVKYPVKDGTANTDVKTHVLIEDMP
GFSYDHEYTVQVKLNAATIKDITGKDVTLKEPVKKDILNTGNVASNNNAGPKLAKPDFDDTNSVQKTASKTEKNAKT
NDSSSMVWYITLFGASFLYLAYRL**GGGGSGGGGS**LTLVVLGVFLFSGWKIGMELYENKHNQTILDDAKAVYTKDAAT
TNVNGEVRDELRDLQKLNKDMVGWLTIIDTEIDYPILQSKDNDYYLHHNYKNEKARAGSIFKDYRNTNEFLDKNTII
YGHNMKDGSMFADLRKYLDKDFLVAHPTFSYESGLTNYEVEIFAVYETTTDFYYIETEFPETTDFEDYLQKVKQQSV
YTSNVKVSGKDRIITLSTCDTEKDYEKGRMVIQGKLVTK
